# *In situ* protein identification and mapping using secondary ion mass spectrometry

**DOI:** 10.1101/803940

**Authors:** Anna M. Kotowska, Philip M. Williams, Jonathan W. Aylott, Alexander G. Shard, Morgan R. Alexander, David J. Scurr

## Abstract

Protein characterisation at surfaces currently requires digestion prior to either liquid extraction of the protein for mass spectrometry analysis or *in situ* matrix-assisted desorption/ionisation. Here, we show that direct assignment of individual proteins and mixtures at surfaces can be achieved by employing secondary ion mass spectrometry (SIMS) with gas cluster ion beam (GCIB) bombardment and an Orbitrap™ analyser. Potential applications of the method are illustrated by demonstrating imaging of a protein film masked by a gold grid and the analysis of a protein monolayer biochip.

## Manuscript

*In situ* analysis of proteins at surfaces, such as in tissue sections^1^, on biomaterials^2^ and patterned biochips^3,4^, is key to understanding the molecular mechanism of disease, developing medicines, biomedical devices and biomarker discovery. Label-free methods of imaging proteins are based on mass spectrometry, with the most widely employed being matrix assisted laser desorption/ionisation (MALDI)^5^. MALDI allows lipids and small peptides containing up to 4 amino acids to be imaged with high lateral resolution (1.4 μm)^6^. However, the analysis of proteins with this technique requires enzymatic digestion and the application of a matrix^7^, which adds a complex preparation step and may modify the structure of the sample and the spatial distribution of analytes in both 2 and 3D. Other notable methods in protein imaging at surfaces are desorption electrospray ionisation (DESI)^8^ and liquid extraction surface analysis mass spectrometry (LESA-MS)^9^. These techniques have limited lateral resolution (>100 μm)^10^ and also require protein digestion prior to sample extraction from the surface.

Time-of-flight secondary ion mass spectrometry (ToF-SIMS) is a surface analysis technique providing label- and matrix-free chemical information of sample surface and in 3D (from a couple of nm to 100s microns into a sample).^11^ Analysis of macromolecules in ToF-SIMS is presently limited by both the sensitivity and the mass resolving power of the ToF analyser. Moreover, proteins are heavily fragmented by energetic primary ion beams, resulting normally in only single amino acid fragments devoid of any primary structural information. Limited information about protein identity, conformation or orientation can be obtained by fingerprint or statistical analysis of amino acid fragment intensities.^12^ This approach has only been demonstrated in simple single-protein samples and cannot be applied to unknown proteins or mixtures.

The emergence of large cluster primary ion source analysis beams, such as C_60_^+^ and the gas cluster ion beam (GCIB) such as Ar_n_^+^ has allowed for more information to be derived from model peptide samples (up to 3kDa) in SIMS by detection of multi amino acid fragments^13^. Nonetheless, proteins have yet to be identified without pre-analysis digestion to peptides in SIMS.

Here, we describe a method of label- and matrix-free identification of intact proteins at surfaces using the 3D OrbiSIMS instrument combining an Ar_3000_^+^ GCIB and an Orbitrap™ analyser^14^. The GCIB reduces fragmentation of the large biomolecules compared to the traditionally employed Bi_3_^+^ liquid metal ion gun (LMIG) as illustrated in Figure 1. The mass accuracy of the Orbitrap™ analyser allows, for the first time, the assignment of characteristic amino acid neutral losses between protein fragments with high certainty (Figure 1b). This method simultaneously allows for relatively high lateral resolution imaging with a high chemical information content shown as insert in Figure 1b and in more detail in Figure S1.

**Figure 1.**
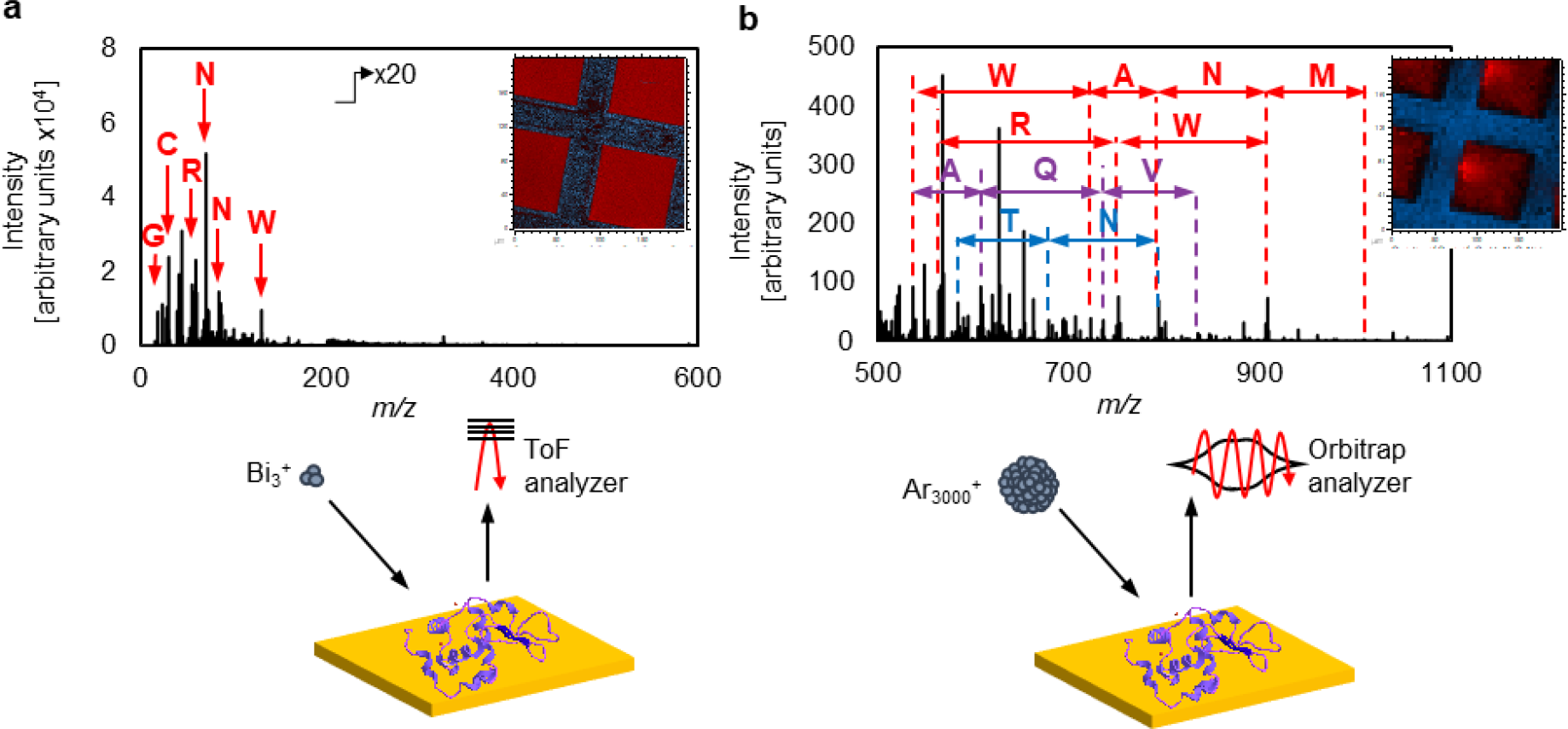
Comparison of (a) Bi_3_+ primary ion beam ToF-SIMS and (b) GCIB Orbitrap™ analysis of lysozyme. In the positive ion secondary ion spectrum obtained using ToF-SIMS, only single amino acid fragments can be identified. In the 3D OrbiSIMS data, protein fragments larger than single amino acids are generated due to the GCIB bombardment. Amino acid neutral losses, W-tryptophan, A-alanine, N-asparagine, M-methionine, R-arginine, Q-glutamine, T-tyrosine, V-valine can be assigned due to the high mass resolving power and mass accuracy the Orbitrap analyser. In the associated ToF-SIMS (a) and 3D OrbiSIMS (b) images of lysozyme film under a gold grid, blue colour represents gold signal and red colour represents a sum of amino acid fragments listed in Table ST25 (a) and assigned peptide fragments listed in Figure S1.

In this study, we prove the concept of using SIMS for direct protein identification from the surface using four example proteins; insulin (~6 kDa)^15^, lysozyme (~14 kDa)^16^, α-chymotrypsin (~50 kDa)^17^ and bovine serum albumin (~133 kDa)^18^. It is immediately apparent that fragments containing multiple amino acids are observed in the spectra (Figure 1b and Figure S4) that can be assigned to specific peptidic secondary ion fragments. The presence of the amino acid neutral losses between peptide fragments indicated the possibility of using *de novo* sequencing to identify the proteins. Continuous sequences of up to 12 amino acid residues were assigned in the spectra of the analysed biomolecules as N-terminal, C-terminal and internal fragments of the sequence (Table ST1). All members of the peptidic sequences can be assigned with confidence due to the high mass accuracy achieved using the Orbitrap™ analyser (<2 ppm) (Figure S2). Further confirmation of the sequencing was performed using the MS/MS capability of the 3D OrbiSIMS instrument to fragment the peptides into their constituent parts (Figure S3). Since the peptidic secondary ion fragments are derived from ballistic fragmentation of the intact protein rather than through fragmentation of enzymatically digested proteins, they cannot be readily compared to existing databases for ‘peptide fingerprint’ identification. However, their utility in identifying the source proteins is explored herein to illustrate the potential for identification of proteins from primary ion-induced protein fragments.

GCIB Orbitrap™ SIMS analysis of the model protein films produced positive ion spectra from which 35% (18 of 51 amino acids), 26% (34 of 129 amino acids), 20% (94 of 482 amino acids) and 5.5% (64 of 1,166 amino acids) of the sequence can be readily assigned from the spectra for insulin, lysozyme, α-chymotrypsin and bovine serum albumin (BSA), respectively (Figure S4 and Table ST1). The sequences were assigned by searching for amino acid neutral losses between the peptidic fragments (*de novo* sequencing approach) predicated on the fragmentation achieved as a result of the primary ion bombardment. Due to the complexity of the protein spectrum generated by the GCIB-induced fragmentation, this sequencing approach is advantageous over the ‘peptide fingerprinting’ method of protein identification used in tandem MS of tryptic peptides. Unlike in MS/MS of peptides, the peptidic ions observed in the positive SIMS spectra are mostly sodium adducts of neutral fragments. Moreover, in addition to b and y ions (normally seen in tandem MS), the spectra obtained with the Ar GCIB contain a, c and z ions (Figure S6, Tables ST2 - ST22). Lastly, in addition to N- and C-terminal ions, a large proportion of the peaks observed in the spectrum are internal fragments of the sequence, not derived from the chain ends (Table ST1). The types of observed peptide fragments are discussed in more detail in the Supplementary Note 1.

Analysis of the human protein sequences listed in the Uniprot database showed that the fraction of proteins which can be unambiguously identified increases with the length of the N-terminal sequence determined after the 3^rd^ residue (Figure S7). In this analysis the first 3 residues were considered by us to be of unknown composition, because experimentally they could not be assigned through sequencing. We found that 7% of the proteins have a unique sequence of residues from 4 to 6, with 63% for residues from 4 to 7, with 89% for residues from 4 to 8 and with 93% for residues from 4 to 9. For the proteins considered, it was possible to generate and readily assign between six and twelve membered peptidic sequences from the N-terminus of the protein. This indicated that *de novo* sequencing of intact proteins in a GCIB Orbitrap™ instrument would be a powerful protein identification method if combined with a spectra database suitable to this method of fragmentation. The potential of this method, as reported herein, to identify proteins is demonstrated in Supplementary Note 2.

An attribute of the 3D OrbiSIMS is the capability to detect picomolar quantities of analyte (Supplementary Note 3). This has been investigated by analysing a protein biochip produced using the method developed by Di Palma *et al.*^19^, which consists of a protein monolayer immobilised on an organic self-assembled monolayer (Figure S9). The reduced amount of analyte allowed for detection of only seven diagnostic peptidic fragments (Table ST24). These peptidic ions were insufficient for sequence assignment, however, by comparison with the reference thin film protein sample the obtained ions were readily identified and assigned.

The ability to image the protein, using the GCIB Orbitrap™, with high chemical specificity and high lateral resolution was investigated by masking a thin protein film (300 nm) using a transmission electron microscopy grid (Figure S1). The secondary ion image of the masked film was achieved with a lateral resolution of 10 μm using the sum of 27 of the characteristic peptidic fragments (Figure S1).

Lastly, identification of proteins in samples containing more than one protein has not previously been achieved in SIMS. We therefore explored the potential of using the *de novo* sequencing method to identify two proteins in a mixture using an equimolar lysozyme and insulin sample. Continuous sequences of up to six amino acids were assigned for both lysozyme and insulin (Figure 2). This analysis of the lysozyme / insulin mixture therefore proved this method to be sufficiently robust to achieve simultaneous identification of both proteins by *de novo* sequencing (Figure S10).

**Figure 2.**
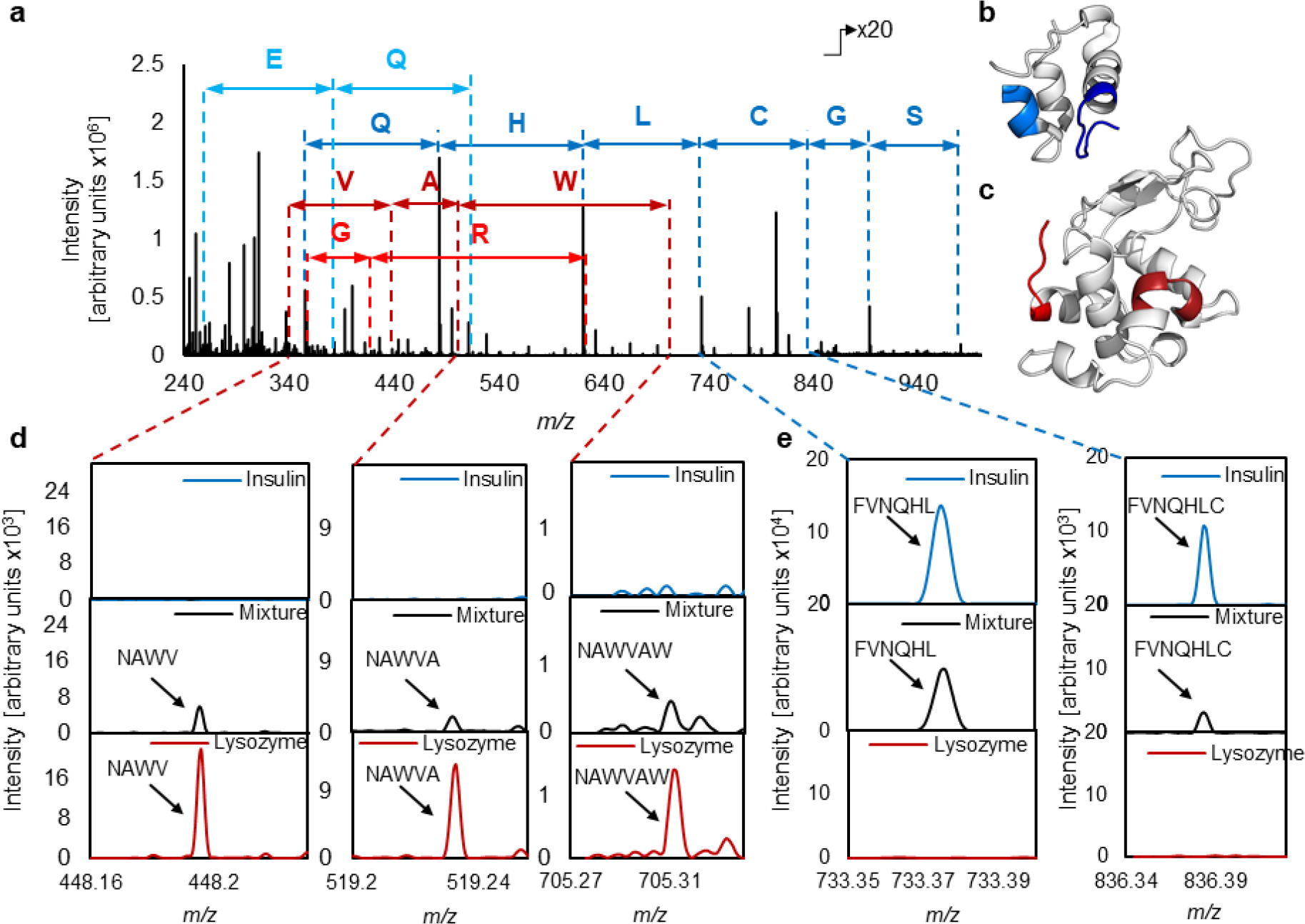
Spectrum of a thin film (300 nm) of 1:1 lysozyme: insulin mixture. Segments of amino acids originating from insulin (light and dark blue) and lysozyme (light and dark red) are presented in the spectrum (a). Segments of insulin sequence observed in the spectrum are highlighted blue in the insulin cartoon (b). Segments of lysozyme sequence observed in the spectrum are highlighted red in the lysozyme cartoon (c). An additional overlay of insulin, mixture and lysozyme samples was magnified to demonstrate lysozyme (d) and insulin (e) related fragments. Peaks assigned as lysozyme sequence NAWVAW (dark red) are detected in the mixture spectrum (black) and are absent in the insulin spectrum (blue) (d). Peaks assigned as insulin sequence FVNQHLC (blue) are detected in the mixture spectrum (black) and are absent in the lysozyme spectrum (red) (e).

In summary, this work demonstrates the first assignment of proteins by *in situ* matrix-free and label-free 3D OrbiSIMS analysis of thin films (300 nm) and immobilised protein monolayers, using a bottom-up *de novo* sequencing method. This analysis is possible without any special sample treatment, such as enzymatic digestion or extraction from the system of interest, instead relying on GCIB fragmentation to produce information-rich peptidic secondary ions and accurate mass assignment with the Orbitrap™ analyser. This approach enables imaging of proteins with high chemical specificity and a lateral resolution of 10 μm using peptidic secondary ion fragments. Additionally, we have demonstrated the first known simultaneous detection and identification of more than one protein within a model mixture using SIMS. The method could in principle be utilised on any instrument with a combined GCIB primary ion beam and analyser with mass resolving power better than 240,000. We anticipate application of this approach in research identifying and imaging protein adsorption on biomaterial surfaces, the development of biosensors reliant upon protein-surface interactions and potentially in tissue sections for the study of disease. It could also be readily adapted to high throughput proteomics analysis of arrayed samples. The potential to apply this method to other aspects of proteomics, such as capturing the native state of the protein using cryogenics, detection of post translational modification or protein-protein interactions or biopolymer analysis (such as DNA, RNA or polysaccharides) is as-yet to be assessed.

## Methods

### Sample preparation

Proteins: lysozyme from chicken egg white, α-chymotrypsin from bovine pancreas type II, insulin solution human, recombinant and bovine serum albumin were purchased from Sigma Aldrich. Hydrogen peroxide (H_2_O_2_) 30% w/w solution, sulfuric acid (H_2_SO_4_) reagent grade 95-95% and N,N-dimethylformamide for molecular biology ≥99% were purchased from Sigma Aldrich. Amicon Ultra 0.5 mL Centrifugal Filters were purchased from Merck Milipore. Insulin, lysozyme and chymotrypsin were purified for the 3D OrbiSIMS analysis. Protein purification was done using Milipore Amicon Ultracentrifuge filter units. The purification was done by following the protocol provided by Milipore: 0.5 mL of 100 μM protein solution in MiliQ water (18 Ω) was placed in the Amicon filter units. The units were centrifuged for 30 minutes at 14,000 g. The protein concentrate was recovered by inverting the filter unit in a clean tube and centrifugation for 2 minutes at 1,000 g. The protein concentration was recovered to 100 μM by refilling the solution with MiliQ water to 0.5 mL. Gold slides were cleaned by UV and sonication in isopropanol for 10 minutes, subsequently rinsed with MiliQ water and dried under Argon flow. Purified proteins were spotted and dried onto a gold slide 3 times to obtain a thick protein film. Protein:peptide mixture was prepared by 1:1 volumetric mixing of 100 μM insulin and 100 μM lysozyme concentrations. The mixture of purified lysozyme and insulin was spotted and dried onto a gold slide 3 times to obtain a thick protein film. Gold grids for electron transmission microscopy (TEM) of 200 mesh × 125 μm pitch were purchased from Sigma Aldrich. For the imaging experiment, the bare gold grid was placed on top of the protein film sample. A bare gold slide and a bare gold grid have been analysed as control samples.

For the preparation of protein monolayers, the gold substrates were cleaned by immersion in piranha solution (70% H_2_SO_4_, 30% H_2_O_2_) at room temperature for 8 minutes, rinsed with MiliQ water and dried with an argon flow. (Caution: Piranha solution reacts violently with organic solvents and should be handled with care).The clean gold substrates were immersed for 24 in 50 μM DMF solutions of heptakis-(6-deoxy-6-thio)-β-cyclodextrin (TCD) for preparation of a self assembled molonayer (SAM). Subsequently, the gold substrates were rinsed with DMF and MiliQ water and dried under an argon flow. The TCD SAMs were immersed in a 0.05 mM lysozyme in PBS solution for 2 h. Following protein immobilization, the samples were washed with PBS buffer followed by submersion in MiliQ water for 1 min. The samples were then dried under argon and placed in the instrument for the analysis immediately.

### Data acquisition

For the acquisition of the 3D OrbiSIMS spectra, a 20 keV Ar_3000_^+^ analysis beam of 20 μm diameter, was used as primary ion beam. Ar_3000_^+^ with duty cycle set to 4.4%, GCIB current was 218 pA. The Q Exactive depth profile was run on the area of 200 × 200 μm using random raster mode with crater size 280 × 280 μm. The cycle time was set to 400 ms. Optimal target potential varied for different samples, oscillating at approximately +68 V. Argon gas flooding was in operation in order to aid charge compensation, pressure in the main chamber was maintained at 9.0 × 10^−7^ bar. The spectra were collected in positive polarity, in mass range 150 - 2250 *m/z*. The injection time was set to 500 ms. Three separate areas were analysed on each sample and each measurement lasted 30 scans, the total ion dose per measurement was 1.63 × 10^11^. Mass resolving power was set to 240,000 at 200 *m/z*.

For the acquisition of the 3D OrbiSIMS MS/MS, a 20 keV Ar_3000_^+^ analysis beam of 20 μm diameter, was used as primary ion beam. Ar_3000_^+^ with duty cycle set to 4.4%, GCIB current was 218 pA. The Q Exactive depth profile was run on the area of 200 × 200 μm using random raster mode with crater size 280 × 280 μm. The cycle time was set to 400 ms. The target potential was set to +68 V. Argon gas flooding was in operation in order to aid charge compensation, pressure in the main chamber was maintained at 9.0 × 10^−7^ bar. The spectra were collected in positive polarity, in mass range 75 - 1125 *m/z*. The injection time was set to 500 ms. Three separate areas were analysed on the sample and each measurement lasted 20 scans, the total ion dose per measurement was 1.63 × 10^11^. Mass resolving power was set to 240,000 at 200 *m/z*. Normalised collision energy (NCE) was set to 35, isolation width was 10 u.

For the acquisition of the 3D OrbiSIMS image, a 20 keV Ar_3000_^+^ analysis beam of 20 μm diameter (imaging of the protein mixture) or a 20 keV Ar_3000_^+^ imaging beam of 5 μm diameter (imaging of lysozyme film masked with a gold grid) was used as primary ion beam. The 20 μm analysis beam was configured as described in the spectra acquisition section. The 5 μm imaging beam duty cycle set to 37.7%, GCIB current was 18 pA. The Q Exactive images were run on the area of 200 × 200 μm using random raster mode. The cycle time was set to 400 ms. Optimal target potential was set to +58 V. Argon gas flooding was in operation in order to aid charge compensation, pressure in the main chamber was maintained at 9.0 × 10^™7^ bar. The images were collected in positive polarity, in mass range 150-2250 *m/z*. The injection time was set to 500 ms. Three separate areas were analysed on each sample and each measurement lasted 1 scan, the total ion dose per measurement was 1.11 × 10^12^. Mass resolving power was set to 240000 at 200 *m/z*.

For the acquisition of LMIG ToF-SIMS image, a 30 keV Bi_3_^+^ primary beam was used. LMIG current was 0.05 pA. The ToF image was run on the area of 200 × 200 μm using random raster mode. The cycle time was set to 250 ms. Optimal target potential was set to +58 V. Three separate areas were analysed on each sample and each measurement lasted 15 scans, the total ion dose per measurement was 9.44 × 10^10^.

The depth of the sputtered material was estimated by the SurfaceLab software as 10 nm per scan and confirmed by profilometry as 300 nm after acquisition of 30 scans. Optical profilometry scans were obtained using a Zeta-20 optical microscope (Zeta Instruments, CA, USA). The scans were acquired in a Z range of 4.6 μm. The number of steps was set to 328, allowing for step size (Z resolution) of 0.014 μm.

The self assembled monolayer and the self assembled monolayer with immobilised lysozyme were measured using an Axis-Ultra XPS instrument (Kratos Analytical, UK) with monochromated Al Kα X-ray source, based at the National Physical Laboratory (NPL), Teddington, UK. Wide scans were acquired for analysis of organic layer thickness on the gold slide with step size 1000 meV, pass energy 160 eV, dwell time 300 ms, in a range 1300 to −10 eV.

The thickness of the self assembled monolayers and t the self assembled monolayer with immobilised lysozyme was measured using J.A. Woollam Co. Inc. alpha-SE™ spectroscopic ellipsometer. The CompleteEASE software was employed to determine the thickness values and the calculations were based on a two-phase organic/Au model, in which the organic layer was assumed to be isotropic and assigned a refractive index of 1.50. The thickness reported is the average of three different measurements on SAM or SAM and protein samples, with the errors reported as standard deviation.

### Data analysis

Thermo Xcalibur and IonToF SurfaceLab 7.1 software were used to process the results and assign the peaks. Xcalibur was used to identify amino acid neutral losses between peaks and SurfaceLab was used to create the peak lists. SurfaceLab was used to measure lateral resolution of ToF-SIMS and 3D OrbiSIMS images. Protein illustrations with highlighted sequences were generated using PyMol™ 2.3.2. Ellipsometry results were analysed using CompleteEASE software. XPS results were analysed using CasaXPS software. Proposed chemical structures of observed protein fragments were produced using ChemDraw Professional 16.0.

