## Supplementary Information for "*In situ* protein identification and mapping using secondary ion mass spectrometry"

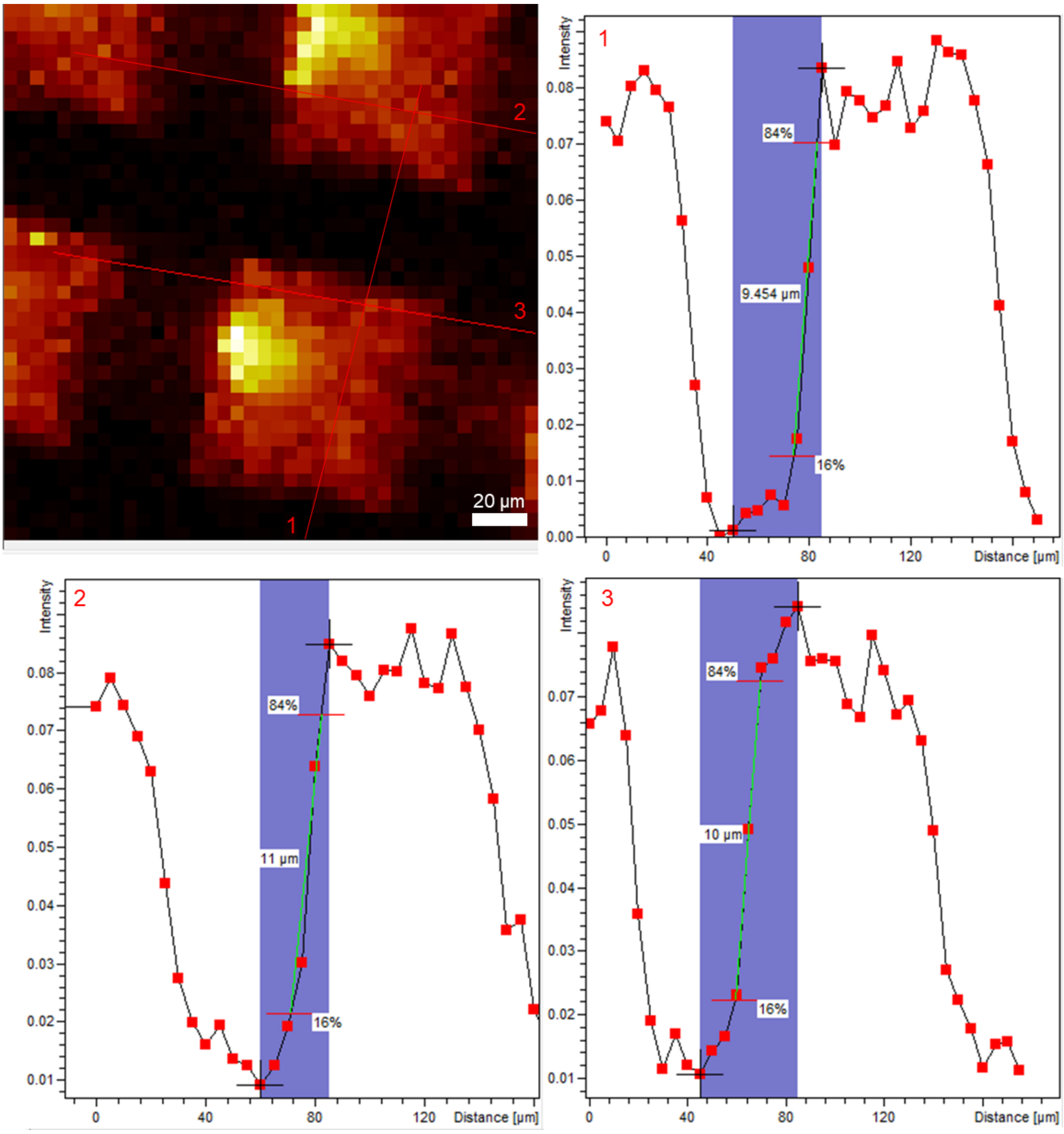


Figure S1 Lateral resolution of the GCIB Orbitrap image obtained with a focussed 5 µm diameter Argon$_{3000}^{+}$ primary beam. The image shows a sum of lysozyme peaks KVFG c, KVFG+Na c, KVFGR a5, KVFGR a5-NH3, KVFGR b5, KVFGR b5-NH3, KVFGR b-CN3H4, KVFGR c, KVFGRC a, KVFGRC b, KVFGRC-S a, KVFGRC-SH2 a, KVFGRCE a, KVFGRCE b, RL y, RL z+1, RL z-1, RG yb, IRG yb, WIRG yb, WIRG ya-NH3, AWIRG yb, AWIRGCRL z-1, VQAWIRG yc, DVQAWIRG yb, NAWV+Na yc, FNTQ+Na a, normalised to total ion count. Line scans 1, 2 and 3 demonstrate the ability to obtain a high chemical specificity image with lateral resolution of 10 µm.

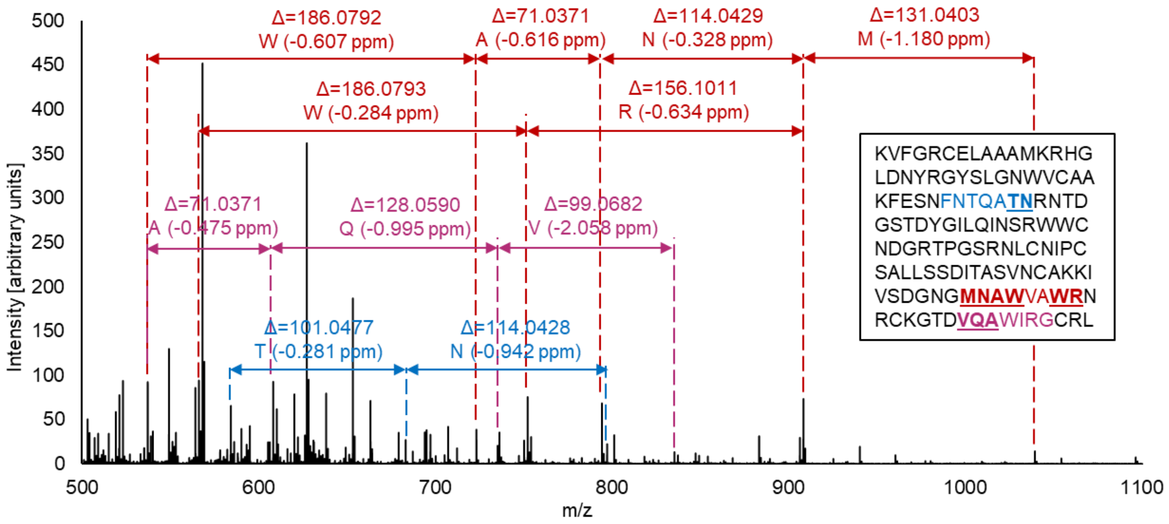


Figure S2 Positive mode GCIB Orbitrap™ spectrum of lysozyme highlighting example sections of amino acid sequence (red, blue, purple), explained by observed peptide fragments. Values in the brackets show deviation of each residue assignment. Amino acid neutral losses can be assigned with confidence due to high mass accuracy of the Orbitrap™ analyser.

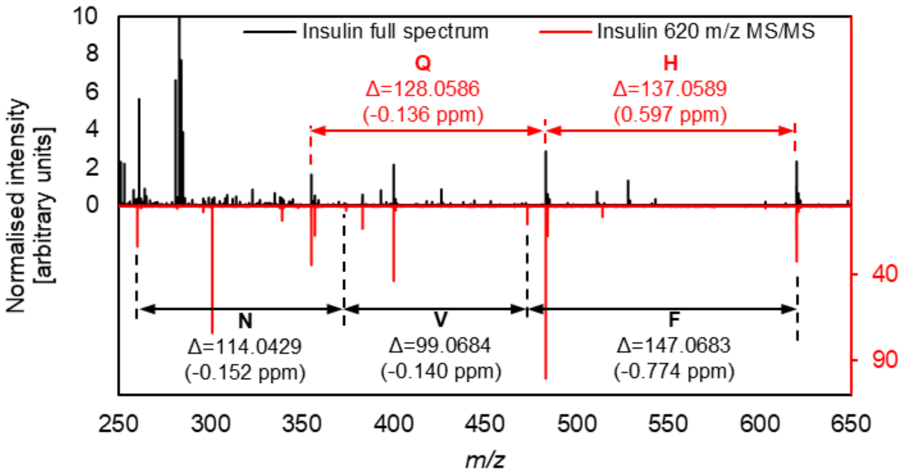


Figure S3 Inverted overlay comparison of full 3D OrbiSIMS spectrum of insulin (black) and 3D OrbiSIMS MS/MS of 620.29 *m/z* (red) ion, assigned as the first 5 amino acids in insulin chain B sequence: FVNQH. All labelled peaks are present in both the full spectrum and the MS/MS spectrum, which confirms the suggested fragmentation. All assigned peaks appear in both full and MS/MS spectra. The peak at 300.05 *m/z* in MS/MS spectrum (red) is an instrument artefact and should be ignored[1].

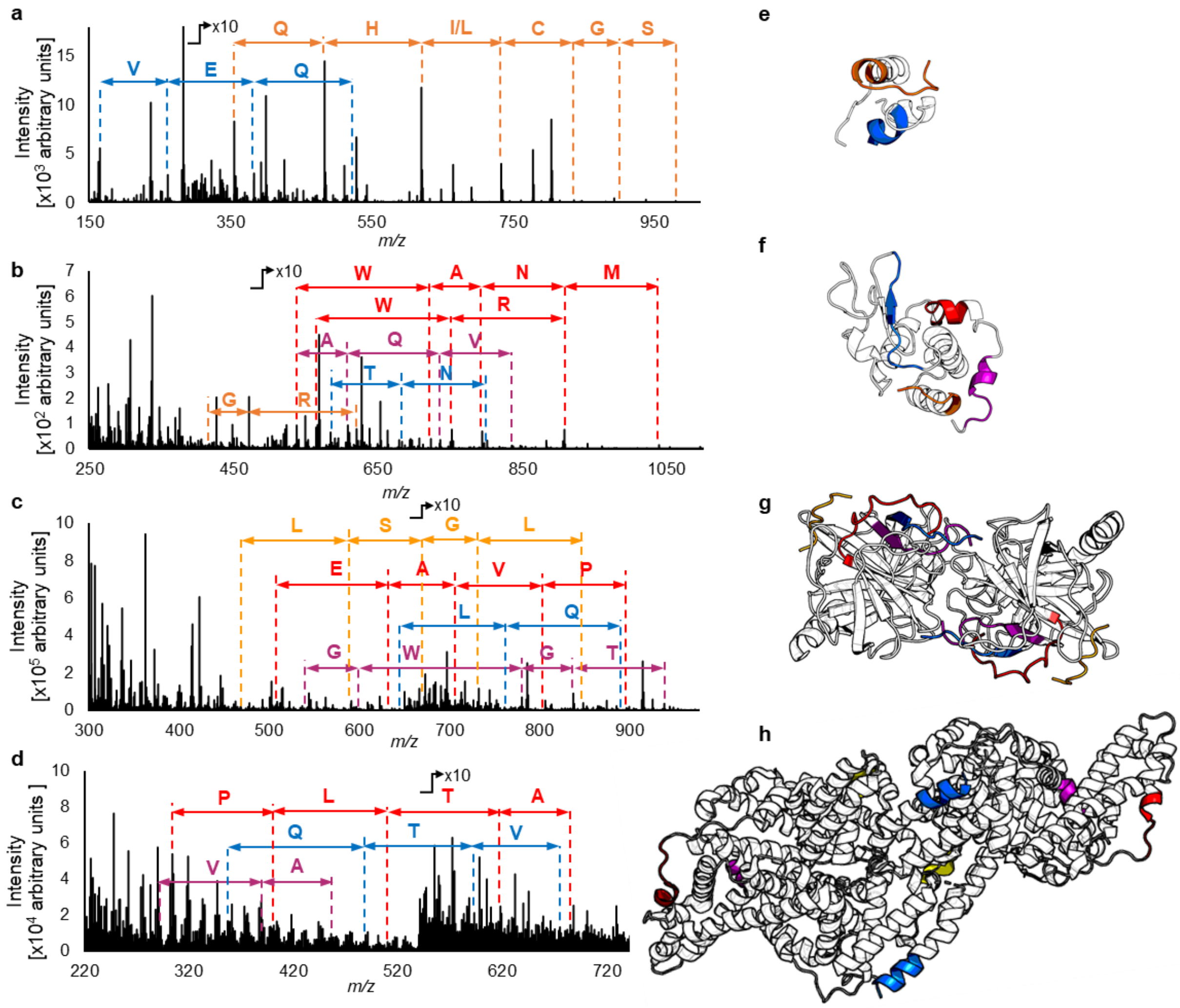

Figure S4 3D OrbiSIMS spectra of insulin (a), lysozyme (b), α-chymotrypsin (c) and BSA (d), with highlighted neutral losses of the amino acids. Colours used to mark amino acid neutral losses in the spectrum correspond to illustrated structures of insulin (e), lysozyme (f), α-chymotrypsin (g) and BSA (h).

**Supplementary Note 1: Types of peptidic fragments generated by Argon GCIB**

The spectra were obtained with a 20 keV $Ar_{3000}^{+}$ primary ion beam, employing ~6.7 eV energy per impact. With such energy, there are the following similarities and differences between commonly used low energy collision induced ionisation (CID). Similarities of observed peptidic fragments with the spectra obtained by (CID) MS/MS of tryptic peptides include presence of a, b and a-$NH_3$ ions[2]. Similar to CID fragmentation, cysteine residues participating in disulphide bonds have been found to form R-S, R-$SH_2$ ions resulting from partial side chain cleavage[3] (Tables ST3, ST5 and ST11). A difference to CID fragmentation is that c fragments, previously seen in electron collision dissociation (ECD) are detected (Figure S6). Each amino acid sequences occurs in the spectra in at least two variants, as both a and c ions (Figure S6b), in most cases as all four a, a-$NH_3$, b and c ions. The presence of internal ions (ya, yb, yc, ya-$NH_3$), not seen in CID fragmentation, is a similarity to high energy collision dissociation (HCD)[4]. The proposed fragment structure of an $Ar_{3000}^{+}$ induced internal fragment ya is presented on an example of NAWVA sequence (Figure S5a). Both y and z ions are detected, z+1 and z+2 ions (Tables ST11, ST15 and ST22), characteristic to MALDI in-source decay (ISD)[5–7] are also observed in our GCIB spectra. The fragment structure of a z+1 ion observed is proposed in an example of RGCRL sequence in Figure S5b.

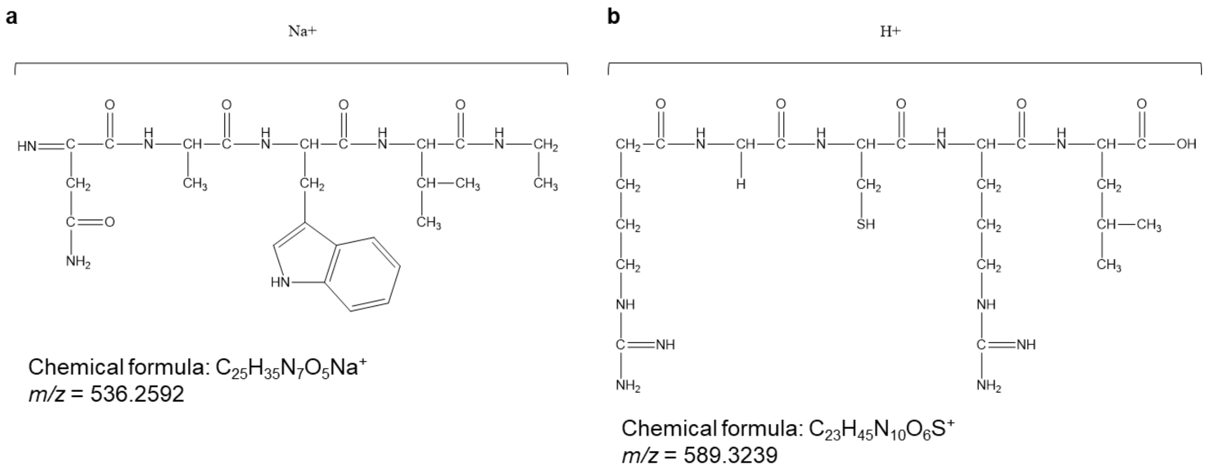


Figure S5. Proposed structures of obtained ions dissimilar to CID MS of tryptic peptides, similar to HCD or MALDI-ISD fragmentation. Internal ion ya is presented on an example of NAWVA sequence observed in the lysozyme spectrum (a). C-terminal ion z+1 is presented on an example of RGCRL sequence observed in the lysozyme spectrum (b). The elemental composition and theoretical *m/z* of the structure was generated in ChemDraw.

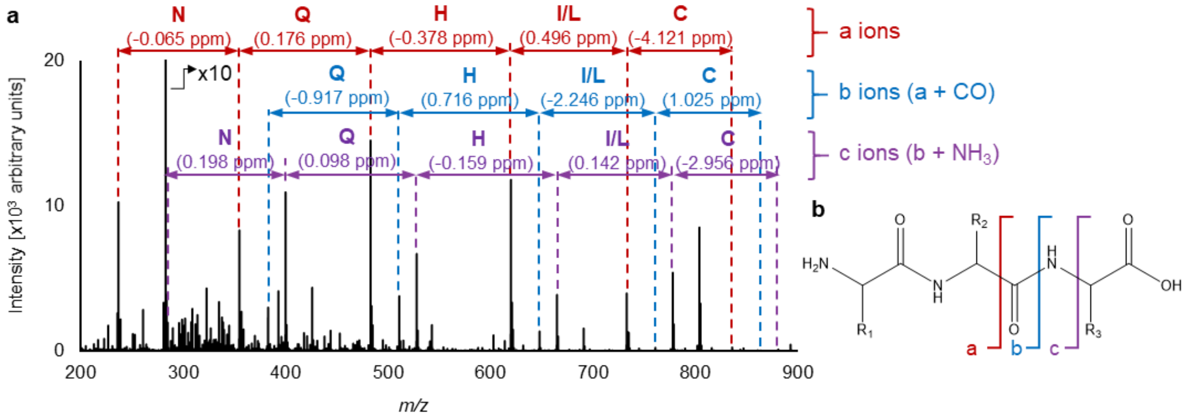


Figure S6 The spectrum (a) shows fragmentation of insulin. The first 8 amino acids of the insulin B chain (FVNQHLC) dominate the spectrum. Each peptide fragment (FVN, FVNQ, FVNQH etc.) is observed as a-, b- and c-type ion. N-terminus ions a, b and c are formed when different bonds related to the peptide bond are broken (b).

Table ST1. Amino acid sequences of human insulin (a), chicken egg lysozyme (b), bovine liver chymotrypsin (c) and bovine serum albumin (d) in FASTA format, as gathered in Uniprot database from 28th July 2019. Parts of sequences, highlighted by colours (blue, orange, yellow, red and purple) represent parts of the amino acid sequence assigned in the spectra. The colours correspond to amino acid segments described in detail in Supplementary Tables 2-21. Bold underlined amino acids are visible in the spectra as a continuous chain of residue neutral losses. The coloured non-underlined sequence is explained by one peak, that is either a di-, tri-, tetra- or pentapeptide or a sodium adduct of those.

**a Human insulin**

>3I40:A|PDBID|CHAIN|SEQUENCE
GI**VEQC**CTSICSLYQLENYCN
>3I40:B|PDBID|CHAIN|SEQUENCE
FV**NQHLCGSHLV**EALYLVCGERGFFYTPKA

**b Lysozyme from chicken egg**

>1AZF:A|PDBID|CHAIN|SEQUENCE
KVF**GRCE**LAAAMKRHGLDNYRGYSLGNWVCAAKFESNFN**TQA**TNRNTDGSTDYGI**LQINS**RWWCNDGRTPGS
RNLCNIPCSALLSSDITASVNCAKKIVSDGNGMNA**WVAWRNR**CKGTD**VQA**WIRGCRL

**c α-chymotrypsin from bovine liver**

>2CHA:A|PDBID|CHAIN|SEQUENCE
**CG**VPAIQPV**LSGL**
>2CHA:B|PDBID|CHAIN|SEQUENCE
IVNGE**EAV**P**GSW**PWQVSLQDKTGFHFCGGSLINENWVVTAAHCGVTTSDVVVAGEFDQGSSSEKIQKLKIAKVF
KNSKYNSLTINNDITLLKLSTAASFSQTVSAVCLPSASDDFAAGTTC**VTTGWGLT**RY
>2CHA:C|PDBID|CHAIN|SEQUENCE
ANTPDR**LQQA**SLPLLSNTNCKKYWGTKIKDAMICAGASGVSSCMGDSGGPLVCKKNGAWTLVGIVSWGSSTCS
TSTPGVYARVTALVNWVQQTLAAN
>2CHA:E|PDBID|CHAIN|SEQUENCE
**CG**VPAIQPV**LSGL**
>2CHA:F|PDBID|CHAIN|SEQUENCE
IVNGE**EAV**PG**SW**PWQVSLQDKTGFHFCGGSLINENWVVTAAHCGVTTSDVVVAGEFDQGSSSEKIQKLKIAKVF
KNSKYNSLTINNDITLLKLSTAASFSQTVSAVCLPSASDDFAAGTTC**VTTGWGLT**RY
>2CHA:G|PDBID|CHAIN|SEQUENCE
ANTPDR**LQQA**SLPLLSNTNCKKYWGTKIKDAMICAGASGVSSCMGDSGGPLVCKKNGAWTLVGIVSWGSSTCS
TSTPGVYARVTALVNWVQQTLAAN

**d Bovine serum albumin**

>3V03:A|PDBID|CHAIN|SEQUENCE
DTH**KSEI**AHRFKDLGEEHFKGLVLIAFSQYLQQCPFDEHVKLVNELTEFAKTCVADESHAGCEKSLHTLFGDELC
KVASLRETYGDMADCCEKQEPERNECFLSHKDDSPDLPKLKPDPNTLCDEFK**ADE**KKFWGKYLYEIARRHPYFY
APELLYYANKYNGVFQECCQAEDKGACLLPKIETMREKVLTSSARQRLRCASIQKFGERAL**KA**WS**VA**RLSQKFP
KAEFVEVTKLVTDLTKVHKECCHGDLLECADDRADLAKYICDNQDTISSKLKECCDKPLLEKSHCIAEVEKDAIP
ENL**PPLTA**DFAEDKDVCKNYQEAKDAFLGSFLYEYSRRHPEYAVSVLLRLAKEYEATLEECCAKDDPHACYSTV
FDKLKHLVDEPQNLIKQNCDQFEKLGEYGFQNALIVRYTRKVPQVSTPTLVEVSRSLGKVGTRCCTKPESERMPC
TEDYLSLILNRLCVLHEKTPVSEKVTKCCTESLVNRRPCFSALTPDETYVPKAFDEKLFTFHADICTLPDTEKQIKK
QTALVELLKHKPKATEEQLKTVMENFVAFVDKCCAADDKEACFAVEGPKL**VVSTQ**TALA
>3V03:B|PDBID|CHAIN|SEQUENCE
DTH**KSEI**AHRFKDLGEEHFKGLVLIAFSQYLQQCPFDEHVKLVNELTEFAKTCVADESHAGCEKSLHTLFGDELC
KVASLRETYGDMADCCEKQEPERNECFLSHKDDSPDLPKLKPDPNTLCDEFK**ADE**KKFWGKYLYEIARRHPYFY
APELLYYANKYNGVFQECCQAEDKGACLLPKIETMREKVLTSSARQRLRCASIQKFGERAL**KA**WS**VA**RLSQKFP
KAEFVEVTKLVTDLTKVHKECCHGDLLECADDRADLAKYICDNQDTISSKLKECCDKPLLEKSHCIAEVEKDAIP
ENL**PPLTA**DFAEDKDVCKNYQEAKDAFLGSFLYEYSRRHPEYAVSVLLRLAKEYEATLEECCAKDDPHACYSTV
FDKLKHLVDEPQNLIKQNCDQFEKLGEYGFQNALIVRYTRKVPQVSTPTLVEVSRSLGKVGTRCCTKPESERMPC
TEDYLSLILNRLCVLHEKTPVSEKVTKCCTESLVNRRPCFSALTPDETYVPKAFDEKLFTFHADICTLPDTEKQIKK
QTALVELLKHKPKATEEQLKTVMENFVAFVDKCCAADDKEACFAVEGPKL**VVSTQ**TALA

Table ST2. Peak list exported from SurfaceLab spectrum of human insulin, consisting of ions detected in the spectrum and assigned as sodium adducts of N-terminal sequence FVNQHLCGSHLV of the B chain of human insulin. The sequence is observed as a, b, c and a-$NH_3$ ions. The *m/z* values represent the experimentally observed center mass of each peak. The deviation (dev.) represents the parts per million (ppm) accuracy of the assignment as distance between the experimental and theoretical *m/z* values. The colour corresponds to the presence of the observed sequence in the whole sequence of human insulin presented in Supplementary Table T1a.

| | **a** | | | **b** | | | **c** | | | **a-$NH_3$** | | |
|---|---|---|---|---|---|---|---|---|---|---|---|---|
| **Description** | ***m/z*** | **Assignment** | **Dev. (ppm)** | ***m/z*** | **Assignment** | **Dev. (ppm)** | ***m/z*** | **Assignment** | **Dev. (ppm)** | ***m/z*** | **Assignment** | **Dev. (ppm)** |
| FVN | 355.1738 | $C_{17}H_{24}N_4O_3Na^+$ | -0.6305 | 383.1688 | $C_{18}H_{24}N_4O_4Na^+$ | -0.5894 | 400.1953 | $C_{18}H_{27}N_5O_4Na^+$ | -0.5872 | 338.1472 | $C_{17}H_{21}N_3O_3Na^+$ | -0.7458 |
| FVNQ | 483.2324 | $C_{22}H_{32}N_6O_5Na^+$ | -0.4139 | 511.2273 | $C_{23}H_{32}N_6O_6Na^+$ | -0.5153 | 528.2539 | $C_{23}H_{35}N_7O_6Na^+$ | -0.4489 | 466.2058 | $C_{22}H_{29}N_5O_5Na^+$ | -0.6667 |
| FVNQH | 620.2912 | $C_{28}H_{39}N_9O_6Na^+$ | -0.4746 | 648.2861 | $C_{29}H_{39}N_9O_7Na^+$ | -0.5111 | 665.3127 | $C_{29}H_{42}N_{10}O_7Na^+$ | -0.4668 | 603.2648 | $C_{28}H_{36}N_8O_6Na^+$ | -0.4038 |
| FVNQHL | 733.3753 | $C_{34}H_{50}N_{10}O_7Na^+$ | -0.3553 | 761.3701 | $C_{35}H_{50}N_{10}O_8Na^+$ | -0.5003 | 778.3967 | $C_{35}H_{53}N_{11}O_8Na^+$ | -0.4391 | 716.3489 | $C_{34}H_{47}N_9O_7Na^+$ | -0.1565 |
| FVNQHLC | 836.3844 | $C_{37}H_{55}N_{11}O_8SNa^+$ | -0.4028 | 864.3798 | $C_{38}H_{55}N_{11}O_9SNa^+$ | 0.0746 | 881.4056 | $C_{38}H_{58}N_{12}O_9SNa^+$ | -0.7268 | x | x | x |
| FVNQHLCG | 893.4058 | $C_{39}H_{58}N_{12}O_9SNa^+$ | -0.4869 | 921.4013 | $C_{40}H_{58}N_{12}O_{10}SNa^+$ | 0.1357 | 938.4272 | $C_{40}H_{61}N_{13}O_{10}SNa^+$ | -0.5241 | x | x | x |
| FVNQHLCGS | 980.4378 | $C_{42}H_{63}N_{13}O_{11}SNa^+$ | -0.4731 | x | x | x | 1,025.4595 | $C_{43}H_{66}N_{14}O_{12}SNa^+$ | -0.2207 | x | x | x |
| FVNQHLCGSH | 1,117.4965 | $C_{48}H_{70}N_{16}O_{12}SNa^+$ | -0.6497 | x | x | x | 1,162.5181 | $C_{49}H_{73}N_{17}O_{13}SNa^+$ | -0.4593 | x | x | x |
| FVNQHLCGSHL | 1,230.5793 | $C_{54}H_{81}N_{17}O_{13}SNa^+$ | -1.6029 | x | x | x | 1,275.5999 | $C_{55}H_{84}N_{18}O_{14}SNa^+$ | -2.1904 | x | x | x |
| FVNQHLCGSHLV | 1,329.6479 | $C_{59}H_{90}N_{18}O_{14}SNa^+$ | -1.3351 | x | x | x | 1,374.6705 | $C_{60}H_{93}N_{19}O_{15}SNa^+$ | -0.4986 | x | x | x |

Table ST3 Peak list exported from SurfaceLab spectrum of human insulin, consisting of ions detected in the spectrum and assigned as N-terminal sequence FVNQHLCGSH of the B chain of human insulin, with sulphur (black letter S) and $SH_2$ removed from the structure. The sequence is observed as a, b and c ions. The *m/z* values represent the experimentally observed center mass of each peak. The deviation (dev.) represents the parts per million (ppm) accuracy of the assignment as distance between the experimental and theoretical *m/z* values. The colour corresponds to the presence of the observed sequence in the whole sequence of human insulin presented in Supplementary Table T1a.

| | **a** | | | **b** | | | **c** | | |
|---|---|---|---|---|---|---|---|---|---|
| **Description** | ***m/z*** | **Assignment** | **Dev. (ppm)** | ***m/z*** | **Assignment** | **Dev. (ppm)** | ***m/z*** | **Assignment** | **Dev. (ppm)** |
| FVNQHLC-S | 804.4124 | $C_{37}H_{55}N_{11}O_8Na^+$ | -0.3555 | 832.4074 | $C_{38}H_{55}N_{11}O_9Na^+$ | -0.24362 | 849.4339 | $C_{38}H_{58}N_{12}O_9Na^+$ | -0.3822 |
| FVNQHLC-$SH_2$ | 802.3967 | $C_{37}H_{53}N_{11}O_8Na^+$ | -0.4309 | 830.3914 | $C_{38}H_{53}N_{11}O_9Na^+$ | -0.66818 | 847.4182 | $C_{38}H_{56}N_{12}O_9Na^+$ | -0.4367 |
| FVNQHLCG-S | 861.4338 | $C_{39}H_{58}N_{12}O_9Na^+$ | -0.4498 | x | x | x | 906.4555 | $C_{40}H_{61}N_{13}O_{10}Na^+$ | -0.1694 |
| FVNQHLCG-$SH_2$ | 859.4181 | $C_{39}H_{56}N_{12}O_9Na^+$ | -0.5272 | x | x | x | 904.4393 | $C_{40}H_{59}N_{13}O_{10}Na^+$ | -0.7691 |
| FVNQHLCGS-S | 948.4657 | $C_{42}H_{63}N_{13}O_{11}Na^+$ | -0.5271 | x | x | x | 993.4865 | $C_{43}H_{66}N_{14}O_{12}Na^+$ | -1.1499 |
| FVNQHLCGS-$SH_2$ | 946.4503 | $C_{42}H_{61}N_{13}O_{11}Na^+$ | -0.3125 | x | x | x | 991.4713 | $C_{43}H_{64}N_{14}O_{12}Na^+$ | -0.7005 |
| FVNQHLCGSH-S | 1,085.5246 | $C_{48}H_{70}N_{16}O_{12}Na^+$ | -0.4644 | x | x | x | x | x | x |
| FVNQHLCGSH-$SH_2$ | 1,083.5094 | $C_{48}H_{68}N_{16}O_{12}Na^+$ | -0.0919 | x | x | x | x | x | x |

Table ST4 Peak list exported from SurfaceLab spectrum of human insulin, consisting of ions detected in the spectrum and assigned as sodium adducts of N-terminal sequence GIVEQC of the B chain of human insulin. The sequence is observed as a, b and c ions. The *m/z* values represent the experimentally observed center mass of each peak. The deviation (dev.) represents the parts per million (ppm) accuracy of the assignment as distance between the experimental and theoretical *m/z* values. The colour corresponds to the presence of the observed sequence in the whole sequence of human insulin presented in Supplementary Table T1a.

| | **a** | | | **b** | | | **c** | | |
|---|---|---|---|---|---|---|---|---|---|
| **Description** | ***m/z*** | **Assignment** | **Dev. (ppm)** | ***m/z*** | **Assignment** | **Dev. (ppm)** | ***m/z*** | **Assignment** | **Dev. (ppm)** |
| GI | 165.0994 | $C_7H_{14}N_2ONa^+$ | -2.3491 | 193.0945 | $C_8H_{14}N_2O_2Na^+$ | -1.2589 | 210.1211 | $C_8H_{17}N_3O_2Na^+$ | -0.7347 |
| GIV | 264.1681 | $C_{12}H_{23}N_3O_2Na^+$ | -0.5409 | 292.1630 | $C_{13}H_{23}N_3O_3Na^+$ | -0.4912 | 309.1895 | $C_{13}H_{26}N_4O_3Na^+$ | -0.6666 |
| GIVE | 393.2106 | $C_{17}H_{30}N_4O_5Na^+$ | -0.7227 | 421.2055 | $C_{18}H_{30}N_4O_6Na^+$ | -0.5668 | 438.2321 | $C_{18}H_{33}N_5O_6Na^+$ | -0.5479 |
| GIVEQ | 521.2692 | $C_{22}H_{38}N_6O_7Na^+$ | -0.5115 | 549.2642 | $C_{23}H_{38}N_6O_8Na^+$ | -0.2508 | x | x | x |
| GIVEQC | 624.2783 | $C_{25}H_{43}N_7O_8SNa^+$ | -0.5621 | x | x | x | x | x | x |

Table ST5 Peak list exported from SurfaceLab spectrum of human insulin, consisting of ions detected in the spectrum and assigned as N-terminal sequence GIVEQC of the B chain of human insulin, with sulphur (black letter S) and $SH_2$ removed from the structure. The sequence is observed as a ions. The *m/z* values represent the experimentally observed center mass of each peak. The deviation (dev.) represents the parts per million (ppm) accuracy of the assignment as distance between the experimental and theoretical *m/z* values. The colour corresponds to the presence of the observed sequence in the whole sequence of human insulin presented in Supplementary Table T1a.

| | **a** | | | **b** | | | **c** | | |
|---|---|---|---|---|---|---|---|---|---|
| **Description** | ***m/z*** | **Assignment** | **Dev. (ppm)** | ***m/z*** | **Assignment** | **Dev. (ppm)** | ***m/z*** | **Assignment** | **Dev. (ppm)** |
| GIVEQC-S | 592.3063 | $C_{25}H_{43}N_7O_8Na^+$ | -0.4222 | x | x | x | x | x | x |
| GIVEQC-$SH_2$ | 590.2913 | $C_{25}H_{41}N_7O_8Na^+$ | 0.6227 | x | x | x | x | x | x |

Table ST6 Peak list exported from SurfaceLab spectrum of chicken egg lysozyme, consisting of ions detected in the spectrum and assigned as sodium adducts of N-terminal sequence KVFGRCE. The sequence is observed as a, b, c and a-$NH_3$ ions. The *m/z* values represent the experimentally observed center mass of each peak. The deviation (dev.) represents the parts per million (ppm) accuracy of the assignment as distance between the experimental and theoretical *m/z* values. The colour corresponds to the presence of the observed sequence in the whole sequence of human insulin presented in Supplementary Table T1b.

| | a | | | b | | | c | | | a-$NH_3$ | | |
|---|---|---|---|---|---|---|---|---|---|---|---|---|
| **Description** | ***m/z*** | **Assignment** | **Dev. (ppm)** | ***m/z*** | **Assignment** | **Dev. (ppm)** | ***m/z*** | **Assignment** | **Dev. (ppm)** | ***m/z*** | **Assignment** | **Dev. (ppm)** |
| KVF | 369.2260 | $C_{19}H_{30}N_4O_2Na^+$ | -0.1771 | 397.2210 | $C_{20}H_{30}N_4O_3Na^+$ | -0.0668 | 414.2473 | $C_{20}H_{33}N_5O_3Na^+$ | -0.5331 | x | x | x |
| KVFG | 426.2475 | $C_{21}H_{33}N_5O_3Na^+$ | -0.1383 | 454.2422 | $C_{22}H_{33}N_5O_4Na^+$ | -0.5043 | 471.2689 | $C_{22}H_{36}N_6O_4Na^+$ | -0.2544 | 409.2207 | $C_{21}H_{30}N_4O_3Na^+$ | -0.7236 |
| KVFGR | 582.3493 | $C_{27}H_{45}N_9O_4Na^+$ | 0.9943 | 610.3447 | $C_{28}H_{45}N_9O_5Na^+$ | 1.7820 | 627.3709 | $C_{28}H_{48}N_{10}O_5Na^+$ | 1.1688 | 565.3217 | $C_{27}H_{42}N_8O_4Na^+$ | -0.7593 |
| KVFGRC | 685.3588 | $C_{30}H_{50}N_{10}O_5SNa^+$ | 1.3660 | 713.3531 | $C_{31}H_{50}N_{10}O_6SNa^+$ | 0.5080 | 730.3795 | $C_{31}H_{53}N_{11}O_6SNa^+$ | 0.2715 | x | x | x |
| KVFGRCE | 814.4001 | $C_{35}H_{57}N_{11}O_8SNa^+$ | -0.3742 | x | x | x | 859.4222 | $C_{36}H_{60}N_{12}O_9SNa^+$ | 0.3087 | x | x | x |

Table ST7 Peak list exported from SurfaceLab spectrum of chicken egg lysozyme, consisting of ions detected in the spectrum and assigned as N-terminal sequence KVFGRCE. The sequence is observed as a, b, c and a-$NH_3$ ions. The *m/z* values represent the experimentally observed center mass of each peak. The deviation (dev.) represents the parts per million (ppm) accuracy of the assignment as distance between the experimental and theoretical *m/z* values. The colour corresponds to the presence of the observed sequence in the whole sequence of human insulin presented in Supplementary Table T1b.

| | a | | | b | | | c | | | a-$NH_3$ | | |
|---|---|---|---|---|---|---|---|---|---|---|---|---|
| **Description** | ***m/z*** | **Assignment** | **Dev. (ppm)** | ***m/z*** | **Assignment** | **Dev. (ppm)** | ***m/z*** | **Assignment** | **Dev. (ppm)** | ***m/z*** | **Assignment** | **Dev. (ppm)** |
| KVFG | x | x | x | x | x | x | 449.2871 | $C_{22}H_{37}N_6O_4^+$ | -0.0082 | x | x | x |
| KVFGR | 560.3669 | $C_{27}H_{46}N_9O_4^+$ | 0.2947 | 588.3618 | $C_{28}H_{46}N_9O_5^+$ | 0.2719 | 605.3884 | $C_{28}H_{49}N_{10}O_5^+$ | 0.3188 | 543.3404 | $C_{27}H_{43}N_8O_4^+$ | 0.38712 |
| KVFGRC | 663.3763 | $C_{30}H_{51}N_{10}O_5S^+$ | 0.5377 | 691.3713 | $C_{31}H_{51}N_{10}O_6S^+$ | 0.7062 | 708.3970 | $C_{31}H_{54}N_{11}O_6S^+$ | -0.4672 | x | x | x |
| KVFGRCE | 792.4187 | $C_{35}H_{58}N_{11}O_8S^+$ | 0.2363 | 820.4125 | $C_{36}H_{58}N_{11}O_9S^+$ | -1.1700 | 837.4403 | $C_{36}H_{61}N_{12}O_9S^+$ | 0.3673 | x | x | x |

Table ST8 Peak list exported from SurfaceLab spectrum of chicken egg lysozyme, consisting of ions detected in the spectrum and assigned as sodium adducts of an internal fragment of the amino acid sequence, GILQINS. The sequence is observed as ya, yb, yc and ya-$NH_3$ ions. The *m/z* values represent the experimentally observed center mass of each peak. The deviation (dev.) represents the parts per million (ppm) accuracy of the assignment as distance between the experimental and theoretical *m/z* values. The colour corresponds to the presence of the observed sequence in the whole sequence of human insulin presented in Supplementary Table T1b.

| | **ya** | | | **yb** | | | **yc** | | | **ya-$NH_3$** | | |
|---|---|---|---|---|---|---|---|---|---|---|---|---|
| **Description** | ***m/z*** | **Assignment** | **Dev. (ppm)** | ***m/z*** | **Assignment** | **Dev. (ppm)** | ***m/z*** | **Assignment** | **Dev. (ppm)** | ***m/z*** | **Assignment** | **Dev. (ppm)** |
| GI | x | x | x | 193.0946 | $C_8H_{14}N_2O_2Na^+$ | -0.7927 | 210.1213 | $C_8H_{17}N_3O_2Na^+$ | -0.2090 | x | x | x |
| GIL | 278.1840 | $C_{13}H_{25}N_3O_2Na^+$ | 0.2682 | 306.1788 | $C_{14}H_{25}N_3O_3Na^+$ | -0.1843 | 323.2053 | $C_{14}H_{28}N_4O_3Na^+$ | -0.1234 | 261.1574 | $C_{13}H_{22}N_2O_2Na^+$ | 0.0814 |
| GILQ | 406.2425 | $C_{18}H_{33}N_5O_4Na^+$ | 0.0092 | 434.2374 | $C_{19}H_{33}N_5O_5Na^+$ | -0.0188 | 451.2644 | $C_{19}H_{36}N_6O_5Na^+$ | 1.0462 | 389.2159 | $C_{18}H_{30}N_4O_4Na^+$ | -0.0761 |
| GILQI | 519.3266 | $C_{24}H_{44}N_6O_5Na^+$ | 0.2001 | 547.3217 | $C_{25}H_{44}N_6O_6Na^+$ | 0.4593 | 564.3480 | $C_{25}H_{47}N_7O_6Na^+$ | 0.0331 | x | x | x |
| GILQIN | 633.3693 | $C_{28}H_{50}N_8O_7Na^+$ | -0.2149 | 661.3646 | $C_{29}H_{50}N_8O_8Na^+$ | 0.3362 | x | x | x | 616.3427 | $C_{28}H_{47}N_7O_7Na^+$ | -0.3422 |
| GILQINS | 720.4005 | $C_{31}H_{55}N_9O_9Na^+$ | -1.3281 | x | x | x | x | x | x | x | x | x |

Table ST9 Peak list exported from SurfaceLab spectrum of chicken egg lysozyme, consisting of ions detected in the spectrum and assigned as sodium adducts of an internal fragment of the amino acid sequence, FNTQA. The sequence is observed as ya, yb, yc and ya-$NH_3$ ions. The *m/z* values represent the experimentally observed center mass of each peak. The deviation (dev.) represents the parts per million (ppm) accuracy of the assignment as distance between the experimental and theoretical *m/z* values. The colour corresponds to the presence of the observed sequence in the whole sequence of human insulin presented in Supplementary Table T1b.

| | **ya** | | | **yb** | | | **yc** | | | **ya-$NH_3$** | | |
|---|---|---|---|---|---|---|---|---|---|---|---|---|
| **Description** | ***m/z*** | **Assignment** | **Dev. (ppm)** | ***m/z*** | **Assignment** | **Dev. (ppm)** | ***m/z*** | **Assignment** | **Dev. (ppm)** | ***m/z*** | **Assignment** | **Dev. (ppm)** |
| FN | 258.1214 | $C_{12}H_{17}N_3O_2Na^+$ | 0.5245 | 286.1163 | $C_{13}H_{17}N_3O_3Na^+$ | 0.3113 | 303.1428 | $C_{13}H_{20}N_4O_3Na^+$ | 0.0698 | x | x | x |
| FNT | 359.1689 | $C_{16}H_{24}N_4O_4Na^+$ | -0.1399 | 387.1639 | $C_{17}H_{24}N_4O_5Na^+$ | -0.0304 | 404.1904 | $C_{17}H_{27}N_5O_5Na^+$ | -0.0471 | 342.1424 | $C_{16}H_{21}N_3O_4Na^+$ | 0.0096 |
| FNTQ | 487.2275 | $C_{21}H_{32}N_6O_6Na^+$ | -0.0114 | 515.2225 | $C_{22}H_{32}N_6O_7Na^+$ | 0.1231 | 532.2500 | $C_{22}H_{35}N_7O_7Na^+$ | 1.8018 | 470.2005 | $C_{21}H_{29}N_5O_6Na^+$ | -1.0115 |
| FNTQA | 558.2651 | $C_{24}H_{37}N_7O_7Na^+$ | 0.8337 | 586.2593 | $C_{25}H_{37}N_7O_8Na^+$ | -0.4676 | 603.2865 | $C_{25}H_{40}N_8O_8Na^+$ | 0.6512 | 541.2381 | $C_{24}H_{34}N_6O_7Na^+$ | -0.0159 |
| NTQA | 411.1963 | $C_{15}H_{28}N_6O_6Na^+$ | 0.2309 | 439.1914 | $C_{16}H_{28}N_6O_7Na^+$ | 0.6327 | x | x | x | 394.1696 | $C_{15}H_{25}N_5O_6Na^+$ | -0.1482 |

Table ST10 Peak list exported from SurfaceLab spectrum of chicken egg lysozyme, consisting of ions detected in the spectrum and assigned as sodium adducts of an internal fragment of the amino acid sequence, NAWVAWRNR. The sequence is observed as ya, yb, yc and ya-$NH_3$ ions. The *m/z* values represent the experimentally observed center mass of each peak. The deviation (dev.) represents the parts per million (ppm) accuracy of the assignment as distance between the experimental and theoretical *m/z* values. The colour corresponds to the presence of the observed sequence in the whole sequence of human insulin presented in Supplementary Table T1b.

| | **ya** | | | **yb** | | | **yc** | | | **ya-$NH_3$** | | |
|---|---|---|---|---|---|---|---|---|---|---|---|---|
| **Description** | ***m/z*** | **Assignment** | **Dev. (ppm)** | ***m/z*** | **Assignment** | **Dev. (ppm)** | ***m/z*** | **Assignment** | **Dev. (ppm)** | ***m/z*** | **Assignment** | **Dev. (ppm)** |
| NA | 180.0743 | $C_6H_{11}N_3O_2Na^+$ | -0.3451 | 208.0692 | $C_7H_{11}N_3O_3Na^+$ | -0.1725 | 225.0958 | $C_7H_{14}N_4O_3Na^+$ | 0.1671 | 163.048 | $C_6H_8N_2O_2Na^+$ | -1.1551 |
| NAW | 366.1534 | $C_{17}H_{21}N_5O_3Na^+$ | -0.7604 | 394.1484 | $C_{18}H_{21}N_5O_4Na^+$ | -0.5582 | 411.1749 | $C_{18}H_{24}N_6O_4Na^+$ | -0.4455 | 349.127 | $C_{17}H_{18}N_4O_3Na^+$ | -0.0065 |
| NAWV | 465.2220 | $C_{22}H_{30}N_6O_4Na^+$ | -0.1686 | 493.2168 | $C_{23}H_{30}N_6O_5Na^+$ | -0.2974 | 510.2430 | $C_{23}H_{33}N_7O_5Na^+$ | -1.1528 | 448.195 | $C_{22}H_{27}N_5O_4Na^+$ | -0.0918 |
| NAWVA | 536.2590 | $C_{25}H_{35}N_7O_5Na^+$ | -0.4148 | 564.2542 | $C_{26}H_{35}N_7O_6Na^+$ | 0.1925 | 581.2808 | $C_{26}H_{38}N_8O_6Na^+$ | 0.2173 | 519.233 | $C_{25}H_{32}N_6O_5Na^+$ | 0.1979 |
| NAWVAW | 722.3385 | $C_{36}H_{45}N_9O_6Na^+$ | -0.0096 | 750.3339 | $C_{37}H_{45}N_9O_7Na^+$ | 0.6554 | 767.3604 | $C_{37}H_{48}N_{10}O_7Na^+$ | 0.6248 | 705.312 | $C_{36}H_{42}N_8O_6Na^+$ | 0.2354 |
| NAWVAWR | 878.4399 | $C_{42}H_{57}N_{13}O_7Na^+$ | 0.3470 | 906.4350 | $C_{43}H_{57}N_{13}O_8Na^+$ | 0.4979 | 923.4617 | $C_{43}H_{60}N_{14}O_8Na^+$ | 0.6619 | x | x | x |
| NAWVAWRN | x | x | x | 1020.4761 | $C_{47}H_{63}N_{15}O_{10}Na^+$ | -1.3684 | x | x | x | x | x | x |
| NAWVAWRNR | x | x | x | 1176.5789 | $C_{53}H_{75}N_{19}O_{11}Na^+$ | 0.2907 | x | x | x | 1,131.5503 | $C_{52}H_{72}N_{18}O_{10}Na^+$ | -6.0393 |

Table ST11 Peak list exported from SurfaceLab spectrum of chicken egg lysozyme, consisting of ions detected in the spectrum and assigned as C-terminal sequence DVQAWIRGCRL. The sequence is observed as y, z-1, z+1 and z+1-$SH_2$ ions. The *m/z* values represent the experimentally observed center mass of each peak. The deviation (dev.) represents the parts per million (ppm) accuracy of the assignment as distance between the experimental and theoretical *m/z* values. The colour corresponds to the presence of the observed sequence in the whole sequence of human insulin presented in Supplementary Table T1b.

| | **y** | | | **z-1** | | | **z+1** | | | **z+1-$SH_2$** | | |
|---|---|---|---|---|---|---|---|---|---|---|---|---|
| **Description** | ***m/z*** | **Assignment** | **Dev. (ppm)** | ***m/z*** | **Assignment** | **Dev. (ppm)** | ***m/z*** | **Assignment** | **Dev. (ppm)** | ***m/z*** | **Assignment** | **Dev. (ppm)** |
| RL | 288.2030 | $C_{12}H_{26}N_5O_3^+$ | -0.0232 | 271.1765 | $C_{12}H_{23}N_4O_3^+$ | -0.0394 | 273.1920 | $C_{12}H_{25}N_4O_3^+$ | -0.2931 | x | x | x |
| CRL | x | x | x | x | x | x | x | x | x | 342.2137 | $C_{15}H_{28}N_5O_4^+$ | 0.4080 |
| GCRL | x | x | x | x | x | x | 433.2231 | $C_{17}H_{33}N_6O_5S^+$ | 0.7444 | x | x | x |
| RGCRL | x | x | x | 587.3074 | $C_{23}H_{43}N_{10}O_6S^+$ | -1.3632 | 589.3241 | $C_{23}H_{45}N_{10}O_6S^+$ | 0.3742 | 555.3364 | $C_{23}H_{43}N_{10}O_6^+$ | 0.4417 |
| IRGCRL | x | x | x | 700.3918 | $C_{29}H_{54}N_{11}O_7S^+$ | -0.7718 | 702.4079 | $C_{29}H_{56}N_{11}O_7S^+$ | -0.0981 | 668.4207 | $C_{29}H_{54}N_{11}O_7^+$ | 0.6856 |
| WIRGCRL | x | x | x | 886.4697 | $C_{40}H_{64}N_{13}O_8S^+$ | -2.1753 | 888.4875 | $C_{40}H_{66}N_{13}O_8S^+$ | 0.2773 | 854.4993 | $C_{40}H_{64}N_{13}O_8^+$ | -0.2662 |
| AWIRGCRL | x | x | x | 957.5085 | $C_{43}H_{69}N_{14}O_9S^+$ | -0.2061 | 959.5243 | $C_{43}H_{71}N_{14}O_9S^+$ | -0.0749 | 925.5357 | $C_{43}H_{69}N_{14}O_9^+$ | -1.0216 |
| QAWIRGCRL | x | x | x | 1085.5692 | $C_{48}H_{77}N_{16}O_{11}S^+$ | 1.7543 | 1087.5831 | $C_{48}H_{79}N_{16}O_{11}S^+$ | 0.1508 | x | x | x |
| VQAWIRGCRL | x | x | x | 1184.6359 | $C_{53}H_{86}N_{17}O_{12}S^+$ | 0.1884 | 1186.6513 | $C_{53}H_{88}N_{17}O_{12}S^+$ | -0.0846 | x | x | x |
| DVQAWIRGCRL | x | x | x | x | x | x | 1301.6790 | $C_{57}H_{93}N_{18}O_{15}S^+$ | 0.5609 | x | x | x |

Table ST12 Peak list exported from SurfaceLab spectrum of α-chymotrypsin from bovine liver, consisting of ions detected in the spectrum and assigned as sodium adducts of internal fragments of the sequence of the A chain of the protein, IQPVLSG. The sequence is observed as ya, yb, yc and ya-$NH_3$ ions. The *m/z* values represent the experimentally observed center mass of each peak. The deviation (dev.) represents the parts per million (ppm) accuracy of the assignment as distance between the experimental and theoretical *m/z* values. The colour corresponds to the presence of the observed sequence in the whole sequence of human insulin presented in Supplementary Table T1c.

| | **ya** | | | **yb** | | | **yc** | | | **ya - $NH_3$** | | |
|---|---|---|---|---|---|---|---|---|---|---|---|---|
| **Description** | ***m/z*** | **Assignment** | **Dev. (ppm)** | ***m/z*** | **Assignment** | **Dev. (ppm)** | ***m/z*** | **Assignment** | **Dev. (ppm)** | ***m/z*** | **Assignment** | **Dev. (ppm)** |
| IQPV | 432.2580 | $C_{20}H_{35}N_5O_4Na^+$ | -0.3296 | 460.2530 | $C_{21}H_{35}N_5O_5Na^+$ | -0.1913 | 477.2796 | $C_{21}H_{38}N_6O_5Na^+$ | 0.0613 | 415.2314 | $C_{20}H_{32}N_4O_4Na^+$ | -0.5261 |
| IQPVL | 545.3421 | $C_{26}H_{46}N_6O_5Na^+$ | -0.0825 | 573.3369 | $C_{27}H_{46}N_6O_6Na^+$ | -0.2980 | 590.3640 | $C_{27}H_{49}N_7O_6Na^+$ | 0.5257 | x | x | x |
| IQPVLS | 632.3745 | $C_{29}H_{51}N_7O_7Na^+$ | 0.5049 | 660.3695 | $C_{30}H_{51}N_7O_8Na^+$ | 0.5169 | x | x | x | x | x | x |
| IQPVLSG | 689.3963 | $C_{31}H_{54}N_8O_8Na^+$ | 0.8434 | 717.3910 | $C_{32}H_{54}N_8O_9Na^+$ | 0.5633 | x | x | x | x | x | x |

Table ST13 Peak list exported from SurfaceLab spectrum of α-chymotrypsin from bovine liver, consisting of ions detected in the spectrum and assigned as C-terminal sequence of the A chain GVPAIQPVLSGL and a full A chain CGVPAIQPVLSGL. The sequence is observed as y ions. The *m/z* values represent the experimentally observed center mass of each peak. The deviation (dev.) represents the parts per million (ppm) accuracy of the assignment as distance between the experimental and theoretical *m/z* values. The colour corresponds to the presence of the observed sequence in the whole sequence of human insulin presented in Supplementary Table T1c.

| **Description** | ***m/z*** | **Assignment** | **Dev. (ppm)** |
|---|---|---|---|
| SGL | 298.1374 | $C_{11}H_{21}N_3O_5Na^+$ | 0.1814 |
| LSGL | 411.2216 | $C_{17}H_{32}N_4O_6Na^+$ | 0.3605 |
| PVLSGL | 607.3429 | $C_{27}H_{48}N_6O_8Na^+$ | 0.4458 |
| IQPVLSGL | 848.4858 | $C_{38}H_{67}N_9O_{11}Na^+$ | 0.6353 |
| PAIQPVLSGL | 1016.5762 | $C_{46}H_{79}N_{11}O_{13}Na^+$ | 1.1239 |
| GVPAIQPVLSGL | 1172.6668 | $C_{53}H_{91}N_{13}O_{15}Na^+$ | 1.5263 |
| CGVPAIQPVLSGL | 1275.6733 | $C_{56}H_{96}N_{14}O_{16}SNa^+$ | -0.6437 |

Table ST14 Peak list exported from SurfaceLab spectrum of α-chymotrypsin from bovine liver, consisting of ions detected in the spectrum and assigned as N-terminal sequence of the B chain IVNGEEAVPGSWPW. The sequence is observed as a, b, and c and ions. The *m/z* values represent the experimentally observed center mass of each peak. The deviation (dev.) represents the parts per million (ppm) accuracy of the assignment as distance between the experimental and theoretical *m/z* values. The colour corresponds to the presence of the observed sequence in the whole sequence of human insulin presented in Supplementary Table T1c.

| | a | | | b | | | c | | |
|---|---|---|---|---|---|---|---|---|---|
| **Description** | ***m/z*** | **Assignment** | **Dev. (ppm)** | ***m/z*** | **Assignment** | **Dev. (ppm)** | ***m/z*** | **Assignment** | **Dev. (ppm)** |
| IVNGE | 507.2538 | $C_{21}H_{36}N_6O_7Na^+$ | 0.1435 | 535.2488 | $C_{22}H_{36}N_6O_8Na^+$ | 0.2665 | 552.2755 | $C_{22}H_{39}N\_{7}O_8Na^+$ | 0.4851 |
| IVNGEE | 636.2966 | $C_{26}H_{43}N_7O_{10}Na^+$ | 0.4378 | 664.2917 | $C_{27}H_{43}N_7O_{11}Na^+$ | 0.6304 | 681.3182 | $C_{27}H_{46}N_8O_{11}Na^+$ | 0.6121 |
| IVNGEEA | 707.3340 | $C_{29}H_{48}N_8O_{11}Na^+$ | 0.8109 | 735.3279 | $C_{30}H_{48}N_8O_{12}Na^+$ | -0.6632 | 752.3554 | $C_{30}H_{51}N_9O_{12}Na^+$ | 0.6360 |
| IVNGEEAV | 806.4025 | $C_{34}H_{57}N_9O_{12}Na^+$ | 0.7118 | 834.3968 | $C_{35}H_{57}N_9O_{13}Na^+$ | -0.0203 | 851.4245 | $C_{35}H_{60}N_{10}O_{13}Na^+$ | 1.3684 |
| IVNGEEAVP | 903.4563 | $C_{39}H_{64}N_{10}O_{13}Na^+$ | 1.8086 | x | x | x | x | x | x |
| IVNGEEAVPG | 960.4762 | $C_{41}H_{67}N_{11}O_{14}Na^+$ | 0.0345 | 988.4701 | $C_{42}H_{67}N_{11}O_{15}Na^+$ | -0.9118 | 1,005.4982 | $C_{42}H_{70}N_{12}O_{15}Na^+$ | 0.6378 |
| IVNGEEAVPGS | 1047.5078 | $C_{44}H_{72}N_{12}O_{16}Na^+$ | -0.3697 | 1075.5042 | $C_{45}H_{72}N_{12}O_{17}Na^+$ | 1.1047 | 1,092.5300 | $C_{45}H_{75}N_{13}O_{17}Na^+$ | 0.3135 |
| IVNGEEAVPGSW | 1233.5870 | $C_{55}H_{82}N_{14}O_{17}Na^+$ | -0.3636 | x | x | x | x | x | x |
| IVNGEEAVPGSWPW | 1516.7212 | $C_{71}H_{99}N_{17}O_{19}Na^+$ | 1.0909 | 1,544.7161 | $C_{72}H_{99}N_{17}O_{20}Na^+$ | 1.0802 | x | x | x |

Table ST15 Peak list exported from SurfaceLab spectrum of α-chymotrypsin from bovine liver, consisting of ions detected in the spectrum and assigned as C-terminal sequence of the B chain IVNGEEAVPGSWPW. The sequence is observed as y, y+Na, and z+1 ions. The *m/z* values represent the experimentally observed center mass of each peak. The deviation (dev.) represents the parts per million (ppm) accuracy of the assignment as distance between the experimental and theoretical *m/z* values. The colour corresponds to the presence of the observed sequence in the whole sequence of human insulin presented in Supplementary Table T1c.

| | **y** | | | **y + Na** | | | **z+1** | | |
|---|---|---|---|---|---|---|---|---|---|
| **Description** | ***m/z*** | **Assignment** | **Dev. (ppm)** | ***m/z*** | **Assignment** | **Dev. (ppm)** | ***m/z*** | **Assignment** | **Dev. (ppm)** |
| RY | 340.1978 | $C_{15}H_{26}N_5O_4^+$ | -0.4456 | 362.1798 | $C_{15}H_{25}N_5O_4Na^+$ | -0.3176 | 323.1712 | $C_{15}H_{23}N_4O_4^+$ | -0.4901 |
| TRY | 441.2455 | $C_{19}H_{33}N_6O_6^+$ | -0.1419 | 463.2279 | $C_{19}H_{32}N_6O_6Na^+$ | 0.6577 | 424.2190 | $C_{19}H_{30}N_5O_6^+$ | -0.2209 |
| LTRY | x | x | x | 576.3118 | $C_{25}H_{43}N_7O_7Na^+$ | 0.3709 | 537.3033 | $C_{25}H_{41}N_6O_7^+$ | 0.2901 |
| GLTRY | x | x | x | x | x | x | 594.3248 | $C_{27}H_{44}N_7O_8^+$ | 0.3256 |
| WGLTRY | x | x | x | x | x | x | 780.4042 | $C_{38}H_{54}N_9O_9^+$ | 0.3274 |
| GWGLTRY | x | x | x | 876.4342 | $C_{40}H_{59}N_{11}O_{10}Na^+$ | 0.3697 | 837.4259 | $C_{40}H_{57}N_{10}O_{10}^+$ | 0.6289 |
| TGWGLTRY | x | x | x | x | x | x | 938.4736 | $C_{44}H_{64}N_{11}O_{12}^+$ | 0.5432 |
| TTGWGLTRY | x | x | x | x | x | x | 1039.5223 | $C_{48}H_{71}N_{12}O_{14}^+$ | 1.5148 |
| VTTGWGLTRY | x | x | x | x | x | x | 1138.5914 | $C_{53}H_{80}N_{13}O_{15}^+$ | 1.9637 |

Table ST16 Peak list exported from SurfaceLab spectrum of α-chymotrypsin from bovine liver, consisting of ions detected in the spectrum and assigned as N-terminal sequence of the C chain ANTPDRLQQA. The sequence is observed as a, b, c and a-$NH_3$ ions. The *m/z* values represent the experimentally observed center mass of each peak. The deviation (dev.) represents the parts per million (ppm) accuracy of the assignment as distance between the experimental and theoretical *m/z* values. The colour corresponds to the presence of the observed sequence in the whole sequence of human insulin presented in Supplementary Table T1c.

| | **a** | | | **b** | | | **c** | | | **a-$NH_3$** | | |
|---|---|---|---|---|---|---|---|---|---|---|---|---|
| **Description** | ***m/z*** | **Assignment** | **Dev. (ppm)** | ***m/z*** | **Assignment** | **Dev. (ppm)** | ***m/z*** | **Assignment** | **Dev. (ppm)** | ***m/z*** | **Assignment** | **Dev. (ppm)** |
| AN | 158.0922 | $C_6H_{12}N_3O_2^+$ | -1.2982 | x | x | x | 203.1137 | $C_7H_{15}N_4O_3^+$ | -0.9171 | x | x | x |
| ANT | x | x | x | 287.1347 | $C_{11}H_{19}N_4O_5^+$ | -0.8970 | x | x | x | 242.1134 | $C_{10}H_{16}N_3O_4^+$ | -0.4241 |
| ANTP | 356.1927 | $C_{15}H_{26}N_5O_5^+$ | -0.4034 | 384.1878 | $C_{16}H_{26}N_5O_6^+$ | 0.0271 | x | x | x | x | x | x |
| ANTPD | 471.2198 | $C_{19}H_{31}N_6O_8^+$ | 0.0197 | x | x | x | x | x | x | x | x | x |
| ANTPDR | 627.3213 | $C_{25}H_{43}N_{10}O_9^+$ | 0.7068 | 655.3164 | $C_{26}H_{43}N_{10}O_{10}^+$ | 0.8687 | 672.3428 | $C_{26}H_{46}N_{11}O_{10}^+$ | 0.7078 | x | x | x |
| ANTPDRL | 740.4056 | $C_{31}H_{54}N_{11}O_{10}^+$ | 0.8150 | 768.4005 | $C_{32}H_{54}N_{11}O_{11}^+$ | 0.8590 | 785.4269 | $C_{32}H_{57}N_{12}O_{11}^+$ | 0.5838 | x | x | x |
| ANTPDRLQ | 868.4644 | $C_{36}H_{62}N_{13}O_{12}^+$ | 0.9576 | 896.4592 | $C_{37}H_{62}N_{13}O_{13}^+$ | 0.8744 | 913.4858 | $C_{37}H_{65}N_{14}O_{13}^+$ | 0.8534 | 851.4392 | $C_{36}H_{59}N_{12}O_{12}^+$ | 2.5788 |
| ANTPDRLQQ | 996.5232 | $C_{41}H_{70}N_{15}O_{14}^+$ | 1.1329 | 1024.5179 | $C_{42}H_{70}N_{15}O_{15}^+$ | 0.8159 | 1041.5444 | $C_{42}H_{73}N_{16}O_{15}^+$ | 0.7937 | 979.4974 | $C_{41}H_{67}N_{14}O_{14}^+$ | 1.9010 |
| ANTPDRLQQA | 1067.5603 | $C_{44}H_{75}N_{16}O_{15}^+$ | 0.9918 | x | x | x | 1112.5812 | $C_{45}H_{78}N_{17}O_{16}^+$ | 0.4736 | 1049.5251 | $C_{44}H_{71}N_{15}O_{15}^+$ | 0.2242 |

Table ST17 Peak list exported from SurfaceLab spectrum of bovine serum albumin, consisting of ions detected in the spectrum and assigned assigned as sodium adducts of N-terminal sequence DTHK. The sequence is observed as a, b and a-$NH_3$ ions. The *m/z* values represent the experimentally observed center mass of each peak. The deviation (dev.) represents the parts per million (ppm) accuracy of the assignment as distance between the experimental and theoretical *m/z* values. The colour corresponds to the presence of the observed sequence in the whole sequence of human insulin presented in Supplementary Table T1d.

| | **a** | | | **b** | | | **c** | | | **a-$NH_3$** | | |
|---|---|---|---|---|---|---|---|---|---|---|---|---|
| **Description** | ***m/z*** | **Assignment** | **Dev. (ppm)** | ***m/z*** | **Assignment** | **Dev. (ppm)** | ***m/z*** | **Assignment** | **Dev. (ppm)** | ***m/z*** | **Assignment** | **Dev. (ppm)** |
| DTH | 348.1278 | $C_{13}H_{19}N_5O_5Na^+$ | -0.1652 | x | x | x | x | x | x | 331.1013 | $C_{13}H_{16}N_4O_5Na^+$ | -0.07559 |
| DTHK | 476.2228 | $C_{19}H_{31}N_7O_6Na^+$ | 0.0452 | 504.218 | $C_{20}H_{31}N_7O_7Na^+$ | 0.4606 | x | x | x | 459.1962 | $C_{19}H_{28}N_6O_6Na^+$ | -0.07392 |

Table ST18 Peak list exported from SurfaceLab spectrum of bovine serum albumin, consisting of ions detected in the spectrum and assigned assigned fragments of N-terminal sequence DTHKSEI. The sequence is observed as a-$NH_3$ ions. The *m/z* values represent the experimentally observed center mass of each peak. The deviation (dev.) represents the parts per million (ppm) accuracy of the assignment as distance between the experimental and theoretical *m/z* values. The colour corresponds to the presence of the observed sequence in the whole sequence of human insulin presented in Supplementary Table T1d.

| | **a** | | | **b** | | | **c** | | | **a-$NH_3$** | | |
|---|---|---|---|---|---|---|---|---|---|---|---|---|
| **Description** | ***m/z*** | **Assignment** | **Dev. (ppm)** | ***m/z*** | **Assignment** | **Dev. (ppm)** | ***m/z*** | **Assignment** | **Dev. (ppm)** | ***m/z*** | **Assignment** | **Dev. (ppm)** |
| DTHK | x | x | x | x | x | x | x | x | x | 437.2155 | $C_{19}H_{29}N_6O_6^+$ | 2.6766 |
| DTHKS | x | x | x | 569.2689 | $C_{23}H_{37}N_8O_9^+$ | 1.9591 | x | x | x | 524.2477 | $C_{22}H_{34}N_7O_8^+$ | 2.5712 |
| DTHKSE | x | x | x | x | x | x | x | x | x | 653.2888 | $C_{27}H_{41}N_8O_{11}^+$ | -0.2240 |
| DTHKSEI | 783.3999 | $C_{33}H_{55}N_{10}O_{12}^+$ | 0.4793 | x | x | x | x | x | x | x | x | x |

Table ST19 Peak list exported from SurfaceLab spectrum of bovine serum albumin, consisting of ions detected in the spectrum and assigned assigned as sodium adducts of internal fragments of the sequence, ADEKKF. The sequence is observed as ya, yb, yc and ya-$NH_3$ ions. The *m/z* values represent the experimentally observed center mass of each peak. The deviation (dev.) represents the parts per million (ppm) accuracy of the assignment as distance between the experimental and theoretical *m/z* values. The colour corresponds to the presence of the observed sequence in the whole sequence of human insulin presented in Supplementary Table T1d.

| | ya | | | yb | | | yc | | | ya-$NH_3$ | | |
|---|---|---|---|---|---|---|---|---|---|---|---|---|
| **Description** | ***m/z*** | **Assignment** | **Dev. (ppm)** | ***m/z*** | **Assignment** | **Dev. (ppm)** | ***m/z*** | **Assignment** | **Dev. (ppm)** | ***m/z*** | **Assignment** | **Dev. (ppm)** |
| KKF | x | x | x | 428.2626 | $C_{21}H_{35}N_5O_3Na^+$ | -1.5439 | x | x | x | x | x | x |
| EKKF | 529.3107 | $C_{25}H_{42}N_6O_5Na^+$ | -0.3328 | 557.3057 | $C_{26}H_{42}N_6O_6Na^+$ | -0.2425 | 574.3321 | $C_{26}H_{45}N_7O_6Na^+$ | -0.3625 | 512.2841 | $C_{25}H_{39}N_5O_5Na^+$ | -0.40883 |
| DEKKF | 644.3376 | $C_{29}H_{47}N_7O_8Na^+$ | -0.3680 | 672.3323 | $C_{30}H_{47}N_7O_9Na^+$ | -0.6363 | 689.3592 | $C_{30}H_{50}N_8O_9Na^+$ | -0.1909 | x | x | x |
| ADEKKF | 715.3753 | $C_{32}H_{52}N_8O_9Na^+$ | 0.4660 | 743.369 | $C_{33}H_{52}N_8O_{10}Na^+$ | -1.1288 | 760.3979 | $C_{33}H_{55}N_9O_{10}Na^+$ | 1.9754 | x | x | x |

Table ST20 Peak list exported from SurfaceLab spectrum of bovine serum albumin, consisting of ions detected in the spectrum and assigned assigned as sodium adducts of internal fragments of the sequence, KAWSVA. The sequence is observed as ya, yb, yc and ya-$NH_3$ ions. The *m/z* values represent the experimentally observed center mass of each peak. The deviation (dev.) represents the parts per million (ppm) accuracy of the assignment as distance between the experimental and theoretical *m/z* values. The colour corresponds to the presence of the observed sequence in the whole sequence of human insulin presented in Supplementary Table T1d.

| | ya | | | yb | | | yc | | | ya-$NH_3$ | | |
|---|---|---|---|---|---|---|---|---|---|---|---|---|
| **Description** | ***m/z*** | **Assignment** | **Dev. (ppm)** | ***m/z*** | **Assignment** | **Dev. (ppm)** | ***m/z*** | **Assignment** | **Dev. (ppm)** | ***m/z*** | **Assignment** | **Dev. (ppm)** |
| WS | 270.1213 | $C_{13}H_{17}N_3O_2Na^+$ | 0.0498 | 298.1162 | $C_{14}H_{17}N_3O_3Na^+$ | 0.0262 | 315.1427 | $C_{14}H_{20}N_4O_3Na^+$ | -0.1394 | 253.0947 | $C_{13}H_{14}N_2O_2Na^+$ | -0.1205 |
| WSV | 369.1895 | $C_{18}H_{26}N_4O_3Na^+$ | -0.4634 | 397.1844 | $C_{19}H_{26}N_4O_4Na^+$ | -0.5907 | 414.2110 | $C_{19}H_{29}N_5O_4Na^+$ | -0.4966 | 352.1631 | $C_{18}H_{23}N_3O_3Na^+$ | -0.2993 |
| WSVA | 440.2265 | $C_{21}H_{31}N_5O_4Na^+$ | -0.7209 | 468.2215 | $C_{22}H_{31}N_5O_5Na^+$ | -0.5507 | 485.2480 | $C_{22}H_{34}N_6O_5Na^+$ | -0.5124 | 423.2001 | $C_{21}H_{28}N_4O_4Na^+$ | -0.3764 |
| AWSVA | 511.2636 | $C_{24}H_{36}N_6O_5Na^+$ | -0.7363 | 539.2586 | $C_{25}H_{36}N_6O_6Na^+$ | -0.4934 | 556.2852 | $C_{25}H_{39}N_7O_6Na^+$ | -0.3611 | 494.2371 | $C_{24}H_{33}N_5O_5Na^+$ | -0.6711 |
| KAWSVA | 639.3583 | $C_{30}H_{48}N_8O_6Na^+$ | -0.8944 | 667.3523 | $C_{31}H_{48}N_8O_7Na^+$ | -2.2938 | 684.3805 | $C_{31}H_{51}N_9O_7Na^+$ | 0.2052 | 622.3320 | $C_{30}H_{45}N_7O_6Na^+$ | -0.5815 |

Table ST21 Peak list exported from SurfaceLab spectrum of bovine serum albumin, consisting of ions detected in the spectrum and assigned assigned as sodium adducts of internal fragments of the sequence, NLPPLTA. The sequence is observed as ya, yb, yc and ya-$NH_3$ ions. The *m/z* values represent the experimentally observed center mass of each peak. The deviation (dev.) represents the parts per million (ppm) accuracy of the assignment as distance between the experimental and theoretical *m/z* values. The colour corresponds to the presence of the observed sequence in the whole sequence of human insulin presented in Supplementary Table T1d.

| | ya | | | yb | | | yc | | | ya-$NH_3$ | | |
|---|---|---|---|---|---|---|---|---|---|---|---|---|
| **Description** | ***m/z*** | **Assignment** | **Dev. (ppm)** | ***m/z*** | **Assignment** | **Dev. (ppm)** | ***m/z*** | **Assignment** | **Dev. (ppm)** | ***m/z*** | **Assignment** | **Dev. (ppm)** |
| NL | 224.1370 | $C_9H_{19}N_3O_2Na^+$ | 0.1761 | 252.1319 | $C_{10}H_{19}N_3O_3Na^+$ | -0.0484 | x | x | x | 207.1104 | $C_9H_{16}N_2O_2Na^+$ | -0.1653 |
| NLP | 321.1897 | $C_{14}H_{26}N_4O_3Na^+$ | -0.1324 | 349.1845 | $C_{15}H_{26}N_4O_4Na^+$ | -0.2465 | 366.2112 | $C_{15}H_{29}N_5O_4Na^+$ | 0.0049 | 304.1632 | $C_{14}H_{23}N_3O_3Na^+$ | -0.0205 |
| NLPP | 418.2423 | $C_{19}H_{33}N_5O_4Na^+$ | -0.3145 | 446.2373 | $C_{20}H_{33}N_5O_5Na^+$ | -0.2253 | 463.2640 | $C_{20}H_{36}N_6O_5Na^+$ | 0.1472 | 401.2158 | $C_{19}H_{30}N_4O_4Na^+$ | -0.3495 |
| NLPPL | 531.3264 | $C_{25}H_{44}N_6O_5Na^+$ | -0.1727 | 559.3213 | $C_{26}H_{44}N_6O_6Na^+$ | -0.2342 | 576.3480 | $C_{26}H_{47}N_7O_6Na^+$ | -0.0113 | 514.2997 | $C_{25}H_{41}N_5O_5Na^+$ | -0.6495 |
| NLPPLT | 632.3741 | $C_{29}H_{51}N_7O_7Na^+$ | -0.1954 | 660.3688 | $C_{30}H_{51}N_7O_8Na^+$ | -0.4899 | 677.3954 | $C_{30}H_{54}N_8O_8Na^+$ | -0.4867 | 615.3473 | $C_{29}H_{48}N_6O_7Na^+$ | -0.5796 |
| NLPPLTA | 703.4108 | $C_{32}H_{56}N_8O_8Na^+$ | -0.8049 | 731.4063 | $C_{33}H_{56}N_8O_9Na^+$ | 0.0712 | x | x | x | 686.3844 | $C_{32}H_{53}N_7O_8Na^+$ | -0.5687 |

Table ST22 Peak list exported from SurfaceLab spectrum of bovine serum albumin, consisting of ions detected in the spectrum and assigned as C-terminal sequence VVSTQTALA. The sequence is observed as y, z and z+2 ions. The *m/z* values represent the experimentally observed center mass of each peak. The deviation (dev.) represents the parts per million (ppm) accuracy of the assignment as distance between the experimental and theoretical *m/z* values. The colour corresponds to the presence of the observed sequence in the whole sequence of human insulin presented in Supplementary Table T1d.

| | **y** | | | **z** | | | **z+2** | | |
|---|---|---|---|---|---|---|---|---|---|
| **Description** | ***m/z*** | **Assignment** | **Dev. (ppm)** | ***m/z*** | **Assignment** | **Dev. (ppm)** | ***m/z*** | **Assignment** | **Dev. (ppm)** |
| TALA | 376.2318 | $C_{16}H_{32}N_4O_6^+$ | 0.3756 | 359.2052 | $C_{16}H_{29}N_3O_6^+$ | 0.3829 | 361.2209 | $C_{16}H_{31}N_3O_6^+$ | 0.4676 |
| QTALA | 506.3067 | $C_{21}H_{42}N_6O_8^+$ | 1.7283 | 489.2795 | $C_{21}H_{39}N_5O_8^+$ | 0.3830 | 491.2950 | $C_{21}H_{41}N_5O_8^+$ | -0.0045 |
| TQTALA | x | x | x | 590.3272 | $C_{25}H_{46}N_6O_{10}^+$ | 0.3040 | 592.3435 | $C_{25}H_{48}N_6O_{10}^+$ | 1.3877 |
| VSTQTALA | x | x | x | x | x | x | 677.3591 | $C_{28}H_{51}N_7O_{12}^+$ | 0.0806 |
| VVSTQTALA | x | x | x | 774.4117 | $C_{33}H_{58}N_8O_{13}^+$ | -0.0820 | x | x | x |

**Supplementary Note 2: Analysis of protein assignment from protein structural database**

Analysis of the Uniprot protein sequences (28th July 2019) and function database was undertaken to determine the degree to which proteins can be identified from sequences deduced from their mass spectrum. For each protein sequence of length n in the database of 21417, all n-l sequences of length l, from 2 to 20 residues, were searched for, with isoleucine substituted for leucine, as they have identical mass. In this analysis first 3 residues were considered by us to be of unknown in composition, because experimentally they could not be assigned through sequencing. The composition of the initial tripeptide can be assigned using the MS/MS capability of the 3D OrbiSIMS instrument. The number of proteins that contained a match for each of its n-l fragment sequences was counted. Figure S7 shows the number of proteins that can be uniquely identified from its sequence from residue 4 to residue 3+l. The total number of proteins which shared at least one sequence with that tested is shown in Figure S8a. The fewest number of proteins that shared a sequence with the test protein is shown in Figure S8b. 89% of the proteins can be identified only by a known N-terminal sequence of 8 amino acids (Figure S7). The capability to readily identify an unknown protein from a N-terminal sequence is limited due to the presence of mid-sequence ions in the spectrum, however the described method provides information about amino acid sequences of sufficient length to enable assignment of 89% of human proteins provided the protein spectra databases are suitably adapted or devised. The method as it is, allows for identification of between 10% (Figure S8a) and 89% (Figure S8b) proteins from a 8-residue sequence found, depending on the composition of the observed sequence. The identification requires following steps:

1. Identify amino acid sequences in the GCIB Orbitrap™ spectrum.

2. Find matching proteins for the longest sequence assuming the sequence originates from a N-terminal.

3. Search for other proteins that share the found sequence in the middle of the amino acid chain. If matches are found, see if any of the other sequences are present in the protein. Choose from proposed proteins ones that are likely to be present in the sample.

4. Go to point 2 until all sequences identified.

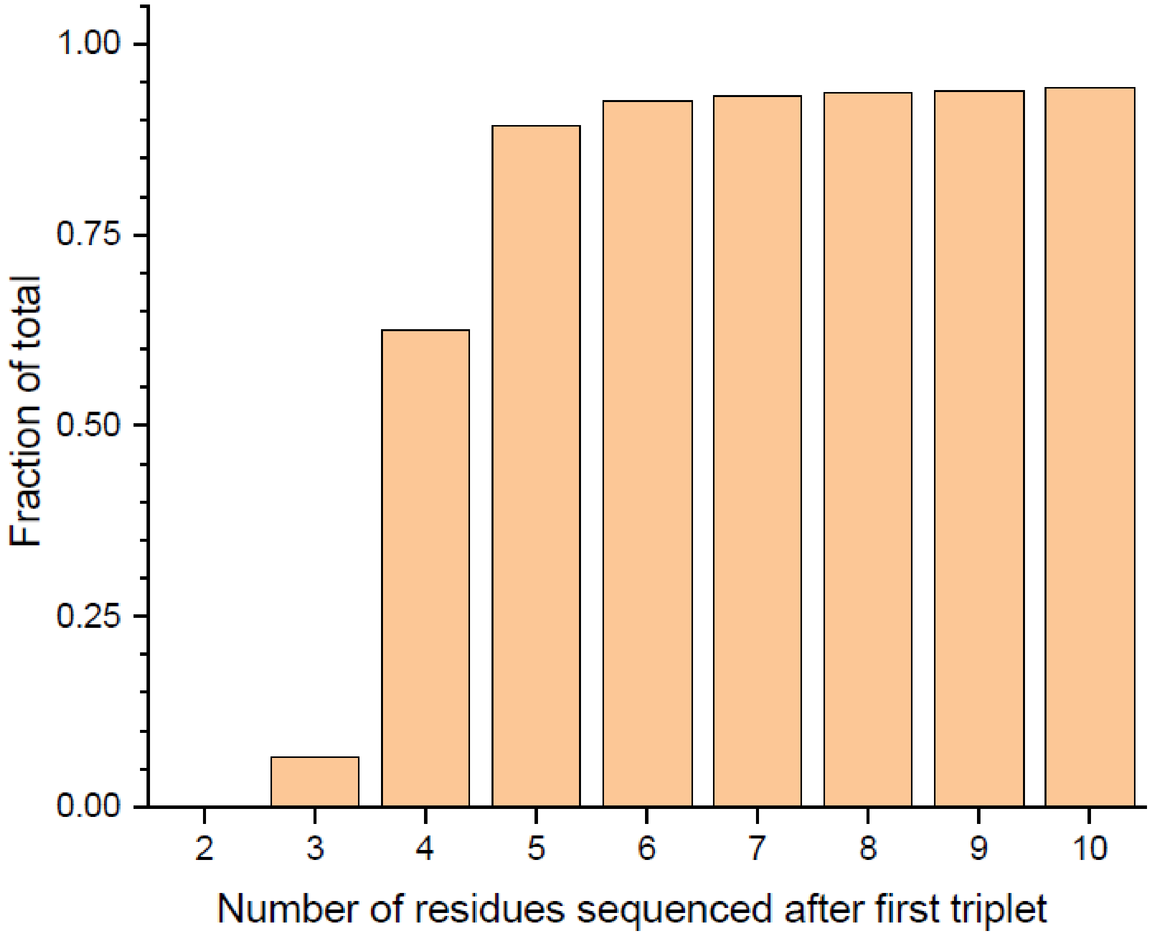


Figure S7 Statistical analysis of the fraction of the proteome that can be identified by N-terminal sequences. 89% of the human proteins in the Uniprot sequence database (28th July 2019) can be confidently identified if the smallest fragment is a tripeptide and is followed by five amino acid residues.

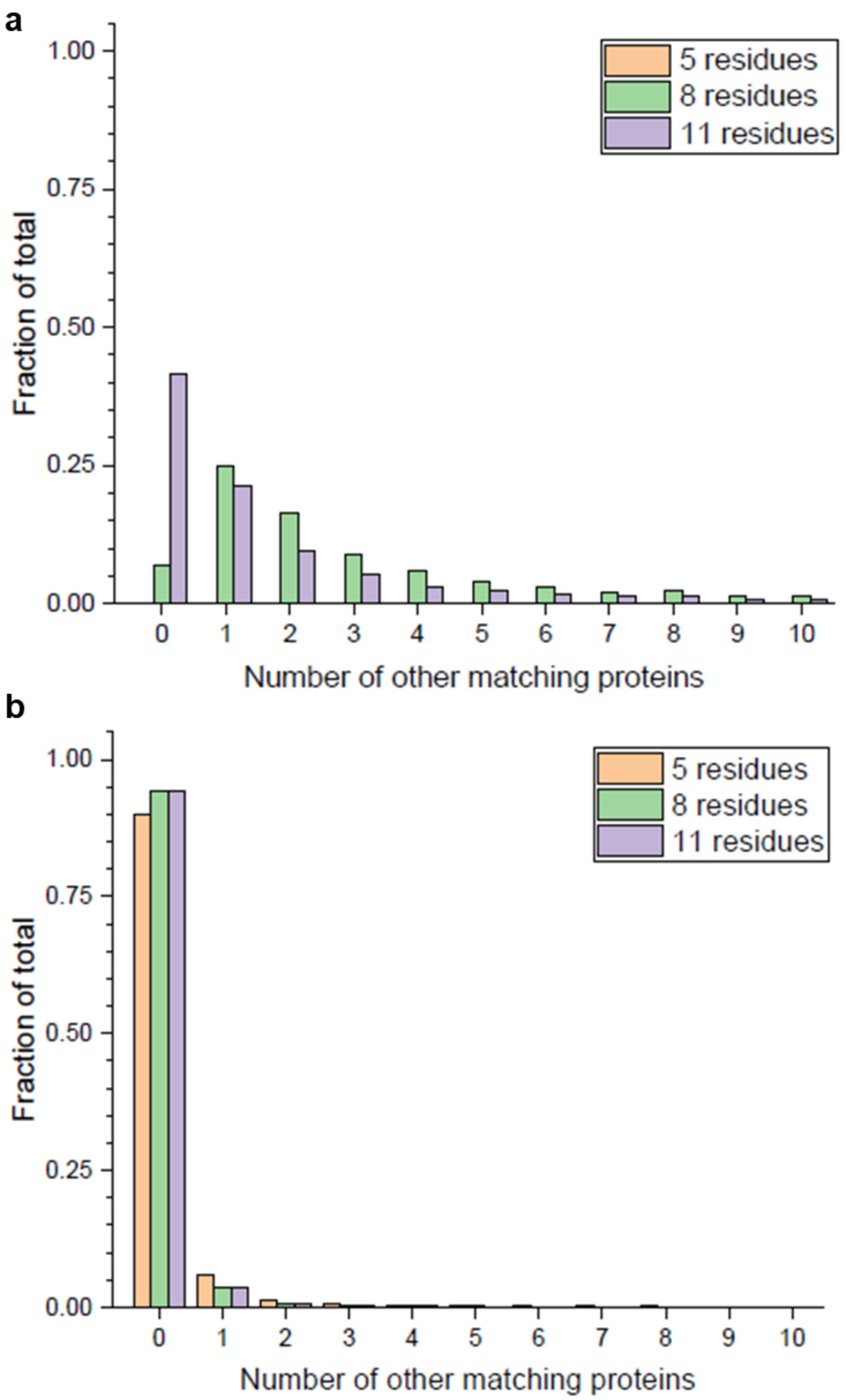


Figure S8 Analysis of human proteins in the Uniprot sequence database (28th July 2019) that share 5, 8 or 11-residue long sequences with any other. Depending on the composition of the sequence found, the sequence may be diagnostic or common to many proteins. (a) 93% of the proteins share an 8-residue long sequence with at least one other protein. (b) 95% of the proteins, however, contain an 8-residue sequence that is unique to the particular protein.

**Supplementary Note 3: Sensitivity and limit of detection of the peptidic fragments**

Sensitivity and limit of detection of the peptidic fragments were calculated for the lysozyme film and the lysozyme monolayer. In lysozyme film samples, the amount of material removed during sputtering is assumed to be pure protein. The amount of protein molecules can be calculated by Equation 1, where $\rho$ is the density of the protein (1.37 g/mL)[8], $A$ is the analysed area, 200 × 200 µm (40000 µm$^2$), $d$ is the material consumed during the analysis, estimated by the SurfaceLab software and confirmed by profilometry (300 nm, Table ST23).

$$n = \frac{\rho A d N_A}{M_W} \qquad \text{Equation 1}$$

In protein monolayer samples, removed material is composed of the protein and the SAM. The amount of protein molecules in the analysed area is calculated by Equation 2. $A$ is the analysed area, 200 × 200 µm (40000 µm$^2$). $A_{TCD}$ is the area of one molecule of thiol-cyclodextrin (TCD) forming the self assembled monolayer.

$$n = \frac{A}{A_{TCD}} \qquad \text{Equation 2}$$

The actual amount of protein molecules is smaller than the amount of cyclodextrin molecules and is determined by the size of the protein, but due to the protein having lost its native structure, the dimensions of the TCD are used in the calculations. The maximum amount of lysozyme molecules in the analysed area is $2.18 \times 10^{10}$ (40 femtomoles). The amount of lysozyme molecules that enabled the protein fragment assignment in the analysed sample was $6.8742 \times 10^{11}$ (1 picomole). The amount of the protein analysed from the monolayer sample (40 femtomoles) is sufficient for the detection of seven diagnostic peaks, however does not enable direct primary structure analysis.

Table ST23 Crater depth measured by optical profilometry. The average of three measurements on one sample (one measurement per crater) is 304 nm. The accuracy (step size) of the profilometer is 14 nm. The standard deviation (SD) of three measueremnts is within the instrument accuracy limit.

| | Cursor Left Avg Ht (µm) | Cursor Right Avg Ht (µm) | Cursor L-R Step (µm) |
|---|---|---|---|
| 1 | 0.3025 | 0.0025 | -0.3 |
| 2 | 0.2976 | 0.0026 | -0.2951 |
| 3 | 0.3398 | 0.0236 | -0.3162 |
| Average | 0.3133 | 0.009567 | -0.30377 |
| SD | 0.02308 | 0.012153 | 0.011043 |

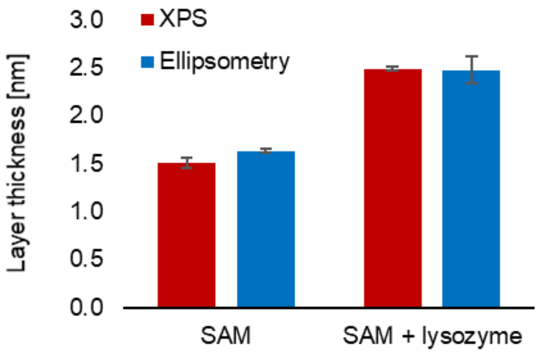


Figure S9 Layer thickness measurements results of the self assembled monolayer (SAM) and SAM with immobilised lysozyme, derived from XPS results (red) and measured by ellipsometry (blue). The error bars represent the standard deviation between three different areas at the surface.

Table ST24 Lysozyme fragments visible in the spectra obtained from a protein monolayer sample. Protein monolayer was obtained by immobilisation of the protein on a self assembled monolayer (SAM) on a gold slide. Three areas on a sample were analysed and a SAM sample without the protein was analysed as a control.

| No | *m/z* | Assignment | Description | SAM + lysozyme | | SAM only | | Bare gold | |
|---|---|---|---|---|---|---|---|---|---|
| | | | | Sample 1 | Sample 2 | Sample 1 | Sample 2 | Sample 1 | Sample 2 |
| 1 | 214.1295 | $C_8H_{16}N_5O_2^+$ | RG | 46277.9 | 59307.63 | 217.33 | 0 | 0 | 0 |
| 2 | 271.1765 | $C_{12}H_{23}N_4O_3^+$ | RL | 3421.62 | 1859.43 | 0 | 0 | 0 | 0 |
| 3 | 449.2864 | $C_{22}H_{37}N_6O_4^+$ | KVFG | 2058.25 | 3253.01 | 0 | 24.75 | 0 | 0 |
| 4 | 560.3665 | $C_{27}H_{46}N_9O_4^+$ | KVFGR | 652.65 | 1236.11 | 0 | 0 | 0 | 0 |
| 5 | 588.3612 | $C_{28}H_{46}N_9O_5^+$ | KVFGR | 9752.09 | 6227.74 | 0 | 0 | 0 | 0 |
| 6 | 605.3875 | $C_{28}H_{49}N_{10}O_5^+$ | KVFGR | 19300.05 | 14136.18 | 0 | 33.58 | 0 | 0 |
| 7 | 631.403 | $C_{30}H_{51}N_{10}O_5^+$ | KVFGRC | 6798.67 | 3267.04 | 0 | 0 | 0 | 0 |

Table ST25 Non-specific amino acid fragments used in ToF-SIMS of proteins, as first assigned by Wagner and Castner [22].

| number | m/z | Assignment | Possible fragment origin |
|---|---|---|---|
| 1 | 30.0351 | $CH_4N^+$ | G |
| 2 | 44.0119 | $CH_2NO^+$ | N |
| 3 | 44.0500 | $C_2H_6N^+$ | A |
| 4 | 44.9775 | $CHS^+$ | C |
| 5 | 56.0520 | $C_3H_6N^+$ | K |
| 6 | 59.0483 | $CH_5N\__3^+$ | R |
| 8 | 61.0099 | $C_2H_5S^+$ | M |
| 9 | 68.0506 | $C_4H_6N^+$ | P |
| 11 | 70.0269 | $C_3H_4NO^+$ | N |
| 12 | 70.0673 | $C_4H_8N^+$ | R |
| 13 | 71.0089 | $C_3H_3O_2^+$ | S |
| 14 | 72.0431 | $C_3H_6NO^+$ | A |
| 15 | 72.0804 | $C_4H_{10}N^+$ | V |
| 16 | 74.0583 | $C_3H_8NO^+$ | T |
| 17 | 76.0199 | $C_2H_6SN^+$ | C |
| 18 | 81.0345 | $C_4H_5N_2^+$ | H |
| 19 | 83.0474 | $C_5H_7O^+$ | V |
| 20 | 84.0397 | $C_4H_6NO^+$ | E/Q |
| 21 | 84.0842 | $C_5H_{10}N^+$ | I, L, K |
| 22 | 86.0988 | $C_5H_{12}N^+$ | I, L |
| 23 | 87.0504 | $C_3H_7N_2O^+$ | N |
| 24 | 88.0379 | $C_3H_6NO_2^+$ | D |
| 25 | 98.0192 | $C_4H_4NO_2^+$ | N |
| 26 | 100.0820 | $C_4H_{10}N_3^+$ | R |
| 27 | 102.0533 | $C_4H_8NO_2^+$ | E |
| 28 | 107.0443 | $C_7H_7O^+$ | T |
| 29 | 110.0746 | $C_5H_8N_3^+$ | H |
| 30 | 120.0783 | $C_8H_{10}N^+$ | F |
| 31 | 130.0562 | $C_9H_8N^+$ | W |
| 32 | 136.0749 | $C_8H_{10}NO^+$ | T |
| 33 | 145.0937 | $C_{10}H_{11}N^+$ | W |
| 34 | 159.0808 | $C_{10}H_{11}N_2^+$ | W |
| 35 | 170.0616 | $C_{11}H_8NO^+$ | W |

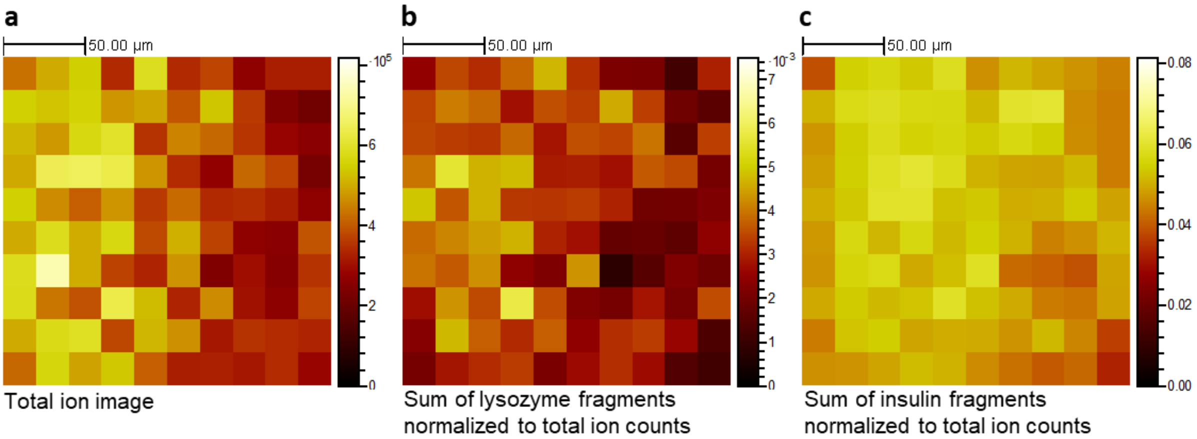


Figure S10 GCIB Orbitrap images of the model mixture The maps represent the total ion image (a), sum of lysozyme fragments, normalized to total (b) and sum of insulin fragments, normalized to total (c). Fragment ions originating from both lysozyme (FN, FNT, FNTQ, FNTQA, NTQA, GI, GIL, GILQ, KVF, KVFG, KVFGR, KVFGRC, KVFGRCE,NA, NAW, NAWV, NAWVA, NAWVAWR, RG) (b) and insulin (FVN, FVNQ, FVNQH, FVNQHL, FVNQHLC, FVNQHLCG, FVNQHLCGS, FVNQHLCGSH, GI, GIV, GIVEQ, GIVEQC) (c) are distributed across the whole analysis area and are detected simultaneously from a mixture.
